# Orthology transfer maps only the conserved core of the *Varroa destructor* proteome and over-calls host absence two times in three

**DOI:** 10.64898/2026.08.20.745999

**Authors:** Štěpán Ryba

## Abstract

The ectoparasitic mite *Varroa destructor* is the principal threat to managed honey bees, and a test case for the genome-scale methods applied to non-model organisms, nearly all of which infer from orthology. We reconstructed the first genome-wide protein-interaction network for *V. destructor* (7,080 proteins, 335,914 interactions), whose modular structure exceeds a degree-preserving null by 368 standard deviations, but whose every edge is interolog-transferred and every node conserved at least to Eukaryota. None of the 791 genes lacking an orthologous group enters it — arithmetic rather than discovery — yet the excluded compartment is large and coherent. It comprises 3,161 genes (30.9% of the proteome), shorter and less annotated than the rest; an annotation-free genome search detects orphans in a tick genome at 4.0% against 70.4% for networked genes. Within the orthology-bearing compartment visibility is non-monotonic: the Acari-level bin (74.1%) falls below the Arthropoda-level bin (89.9%). The same logic applied to host comparison yields a benchmarked error: of genes called absent from *Apis* on group identity alone, 67.4% recover a sequence homologue — against zero for a shuffled null and 1.3% in the presence direction — rising to 78.3% in the least panel-biased stratum. Both figures are properties of the calling rule: under an identity floor the error directions cross near 34% identity; orthology cannot be said to err in either direction without fixing the criterion first. Host divergence resolves into gene absence and residue-level substitution, falling in those two compartments respectively. A bee-sparing target map follows as broader impact.

**Significance statement:** Biologists routinely work out what an unstudied organism’s genes do, and which of them differ from those of its host, by matching those genes to counterparts in well-studied species: a shortcut that is fast, standard and almost never checked. Using the honey-bee parasite *Varroa destructor*, we show that this shortcut has two blind spots that pull in the same direction, because a network built this way contains only the mite’s ancient shared machinery and none of the fast-evolving secreted proteins that make it a parasite, while the same matching, used to decide that a gene is missing from the bee, is wrong about two times in three. Since both errors fall precisely on the genes that distinguish a parasite from its host, comparative work on any non-model organism needs a second, sequence-level check before an absence can be believed.

**Graphical abstract.:** Submitted as a separate file, 4:3 landscape, 160 × 120 mm, TIF at 600 dpi. Not a numbered display item.

## Introduction

*Varroa destructor* (Acari: Mesostigmata) is an obligate ectoparasite of honey bees and the principal biotic threat to managed *Apis mellifera* worldwide. It feeds on developing brood and on adults, vectors a suite of debilitating viruses, and drives the colony losses recorded across a now near-global range (Techer et al. 2019). Two features make it an unusually informative system for asking how a parasite’s proteome is organised. It is a recently host-shifted parasite with a compact, well-assembled genome and no experimental interaction data of its own, which is the situation of most organisms to which genome-scale inference is applied. And half a century of acaricide use has concentrated attention on a small, defined set of neuro-active and detoxification targets (Guo et al. 2021; Inak et al. 2025), giving an unusually sharp external reference against which a genome-wide inventory can be checked.

The standard route to such an inventory is the protein–protein interaction network: hubs tend to be essential, information-flow bottlenecks tend to be regulatory, and modules correspond to pathways and complexes. For a non-model organism no experimental interactions exist, and a network can be obtained only by orthology — “interolog” — transfer, importing interactions from orthologues in better-studied species. STRING implements this at genome scale (Szklarczyk et al. 2023): submitting a proteome returns an entirely prediction-based network built from a chosen reference clade. The approach gives broad coverage and underlies a growing number of non-model interactomes, but it carries a structural property that is rarely stated and, to our knowledge, has not been measured: it can represent only what an organism *shares* with the species from which interactions are transferred. The biology that makes a parasite a parasite — rapidly evolving, lineage-restricted, frequently secreted proteins at the host interface — accrues no transferable orthology, and is therefore excluded by construction rather than by evidence.

Stated that way the exclusion is a tautology, and the interesting questions are the ones that remain once the tautology is set aside: how large the excluded compartment is, whether it is a coherent biological object or a residue of failed annotation, and whether the same orthology-based reasoning also misleads where nothing is definitional. One such place is the comparative call on which most parasite-versus-host claims ultimately rest. Declaring a gene absent from the host because its orthologous-group identifier does not appear in the host proteome is the default output of a group-level comparative join, and it is used to nominate host-specific genes, gene losses and selective targets. Its error rate has not been benchmarked directly, nor has it been asked whether that error is symmetric between the presence and the absence direction, or how much of it is contributed by the composition of the comparison panel rather than by the biology of the organism.

Here we address both questions on a single dataset. We reconstruct the first genome-wide interactome for *V. destructor*, test its modular structure against a degree-preserving null, and characterise what it omits using gene-model, genomic-placement and annotation-free evidence independent of the orthology assignment that defines the omission. We benchmark orthology-based host-absence against a sequence-level search with a shuffled-sequence control, stratify the error by panel support, protein length and annotation status, and repeat the test on the acarine side of the panel using no annotation at all. The results resolve the proteome into two evolutionarily distinct compartments, resolve divergence from the host into two mechanisms — gene absence and residue-level substitution — that fall in those two compartments respectively, and give a quantitative, asymmetric and panel-dependent error rate for the comparative shortcut on which much non-model comparative genomics depends. This study makes no test of selection: its data layer is a single reference proteome, an interolog network and a comparative orthology join. The applied corollary — a route to bee-sparing selectivity — follows from the architecture rather than driving it, and is presented as broader impact.

## Results

### 1. A genome-wide interolog interactome captures the conserved cellular core

#### A single, well-mapped functional network

Submitting the 10,241-gene representative proteome to STRING v12.0 yielded, at the medium-confidence threshold (combined_score ≥ 400), a network of **7,080 proteins and 335,914 unique undirected interactions**, organised into one dominant connected component (7,062 proteins) plus six small peripheral components (Figure 1A). STRING distributes each pair as two directed rows, so this network corresponds to 671,828 rows in the source file; every edge count reported in this study is a count of unique undirected pairs unless stated otherwise. The degree distribution is strongly right-skewed (median 60, mean 94.9, maximum 863), and every node maps unambiguously to a *Varroa* gene with scaffold coordinates. Raising the threshold to combined_score ≥ 700 retained the same architecture more sparsely (5,702 proteins; 92,651 edges). This is, to our knowledge, the first genome-wide protein-interaction network reconstructed for *V. destructor*.

**Fig. 1.**
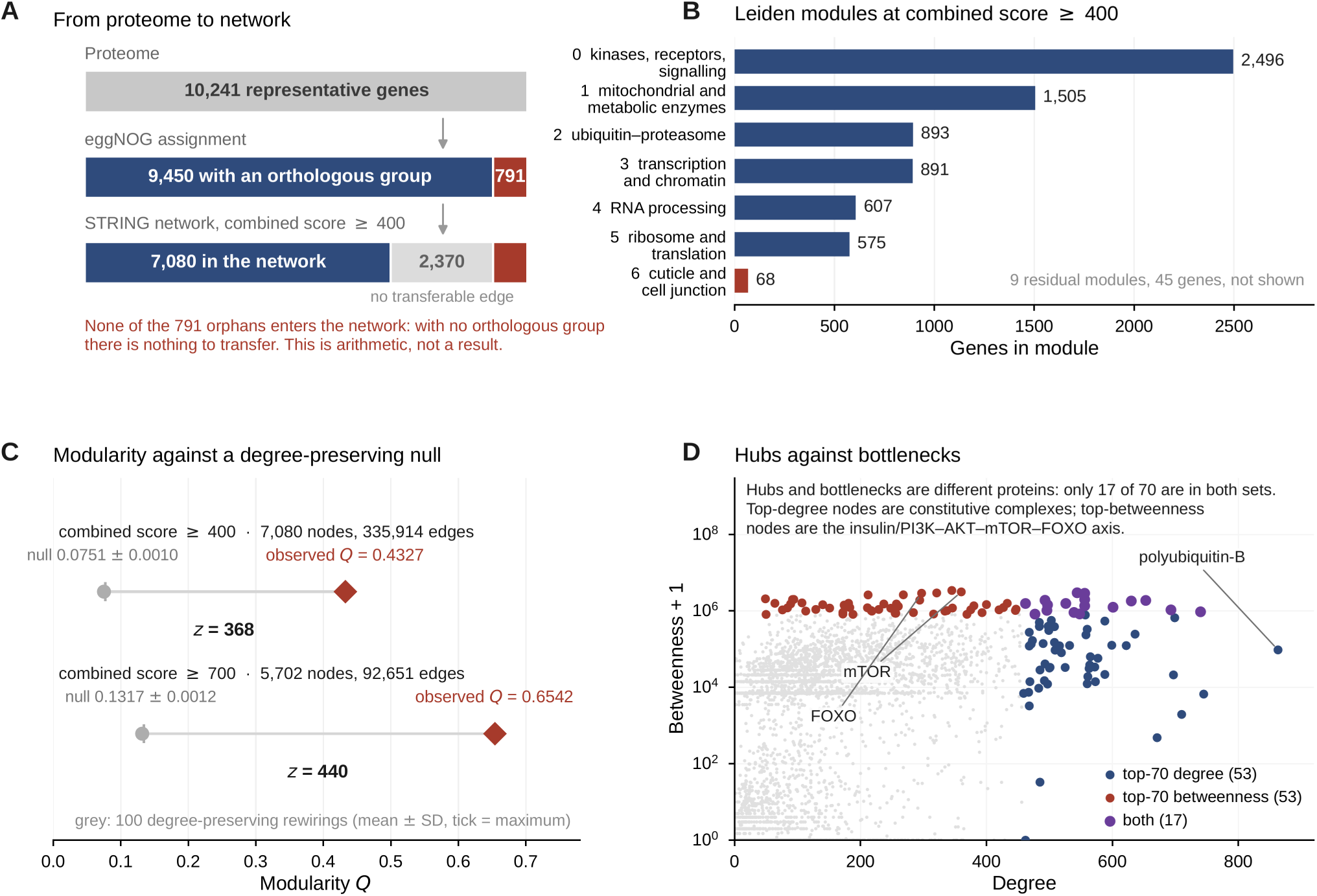
*The* Varroa destructor *interactome is modular beyond its degree sequence, and entirely transferred.* (A) From proteome to network: of 10,241 representative genes, 9,450 carry an eggNOG orthologous group and 791 do not; 7,080 enter the network at combined score ≥ 400, and none of the 791 orphans does, which is arithmetic rather than a result. (B) Leiden module sizes for the seven major modules, from kinases, receptors and signalling (2,496 genes) to a cuticle and cell-junction module (68); nine residual modules holding 45 genes are not shown. Module labels are descriptive summaries of the RefSeq product names of their members. (C) Observed modularity against 100 degree-preserving rewirings: Q = 0.4327 against a null of 0.0751 ± 0.0010 with a maximum of 0.0771 at ≥ 400 (z = 368), and Q = 0.6542 against 0.1317 ± 0.0012 with a maximum of 0.1342 at ≥ 700 (z = 441). (D) Degree against betweenness for all 7,080 nodes. Hubs and bottlenecks are largely different proteins: only 17 of the top 70 by degree are also in the top 70 by betweenness, the top-degree set is dominated by the ribosome and translation module (30 of 70) and the top-betweenness set by the kinase and signalling module (27 of 70), whose highest-ranking members are components of the insulin/PI3K–AKT– mTOR–FOXO axis. Every edge in this network is interolog-transferred; none rests on direct experimental evidence in this organism. *Alt text:* Four-panel figure. A three-step bar showing 10,241 genes narrowing to 7,080 networked with 791 orphans excluded; a horizontal bar chart of seven module sizes; a plot of observed modularity far to the right of a rewired null at two confidence thresholds; and a scatter of degree against betweenness in which the highest-degree and highest-betweenness proteins form two largely separate groups.

#### Modular architecture, tested against a degree-preserving null

Leiden community detection resolved the ≥ 400 network into **16 modules (modularity Q = 0.4327)**, seven of them large. Functional over-representation gave each a clear identity: membrane transport and signalling (M0, n = 2,496), core metabolism and oxidative phosphorylation (M1, 1,505), cell cycle and the ubiquitin–proteasome system (M2, 893), transcription and chromatin (M3, 891), pre-mRNA splicing (M4, 607), ribosome biogenesis and rRNA processing (M5, 575), and a cuticle and cell-junction module (M6, 68; Figure 1B). The smallest module is the one functional class that Figure 2C shows to be least visible: of the cuticular and structural proteins that do enter the network, those that remain form their own module rather than joining the conserved core.

**Fig. 2.**
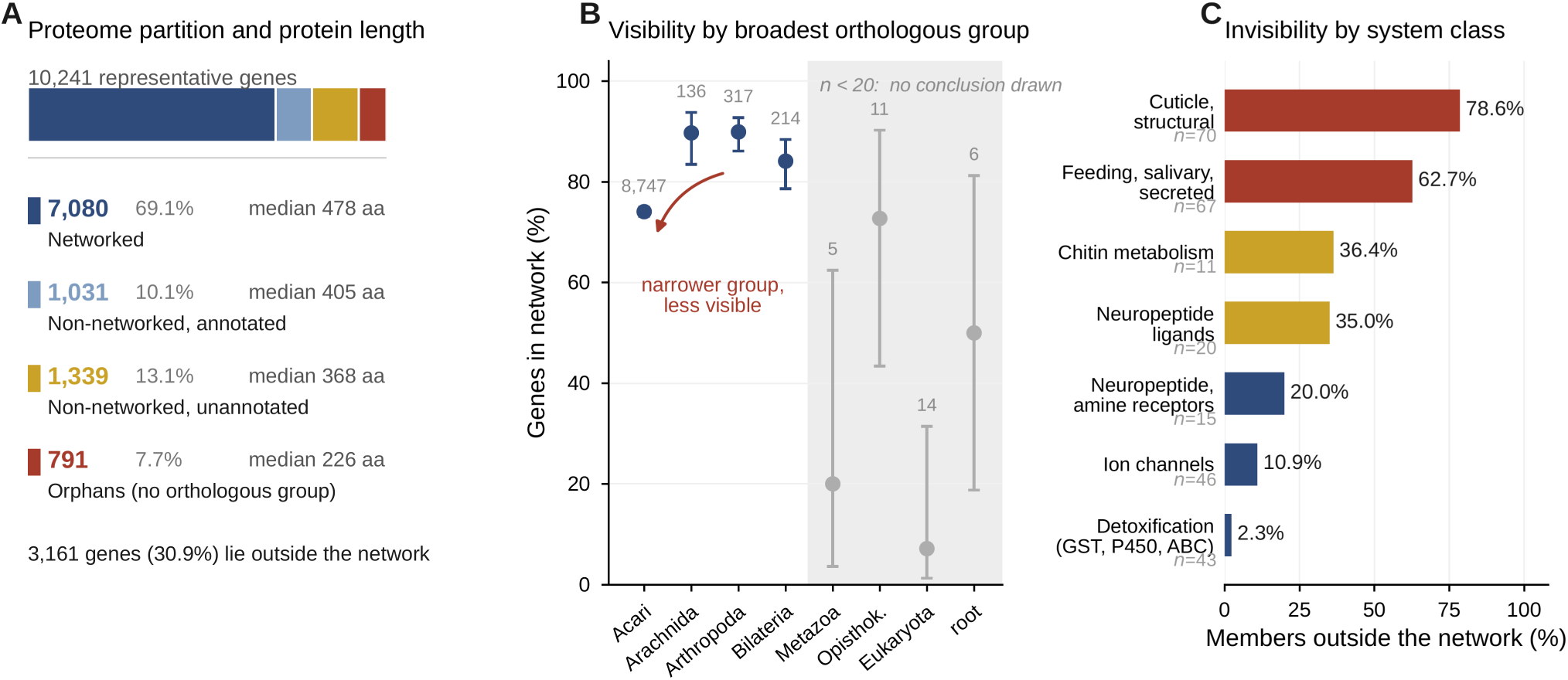
A large, distinct compartment of the proteome lies outside the network, and visibility within the orthology compartment is non-monotonic. (A) Partition of the proteome into four compartments with their median protein lengths: 7,080 networked (478 aa), 1,031 non-networked but functionally annotated (405 aa), 1,339 non-networked and unannotated (368 aa) and 791 orphans (226 aa); 3,161 genes (30.9%) lie outside the network. (B) Percentage of genes networked by the narrowest eggNOG conservation level, with 95% confidence intervals. Among the four bins with n ≥ 20, visibility spans 15.8 percentage points and runs against the expected direction: the Acari-level bin (74.1%, 6,480 of 8,747) falls below the Arthropoda-level bin (89.9%, 285 of 317) with non-overlapping intervals. The four bins with fewer than twenty genes are shown in grey and no conclusion is drawn from them. The ordering is not a length effect — the least visible bin carries the longest median protein, 462 aa against 391 aa — and it holds within both annotation strata, most sharply among unannotated genes (Acari 35.4% against Arthropoda 78.0%). (C) Percentage of annotated members lying outside the network by functional system class, from cuticle and structural proteins (78.6%) to detoxification enzymes (2.3%). Annotation status is the dominant covariate of network membership (odds ratio 11.24) and the class effects survive conditioning on annotation and length. *Alt text:* Three-panel figure. A stacked bar partitioning the proteome into four compartments with median lengths falling from left to right; a point-and-interval plot of percentage networked across conservation levels in which the Acari bin sits below the Arthropoda bin, with small bins greyed; and a horizontal bar chart ranking seven functional classes by the fraction of members outside the network.

This modular structure is not a property of the degree sequence (Figure 1C). Against 100 degree-preserving rewirings of the same graph (10 swap attempts per edge), observed modularity exceeds the randomised expectation by 368 standard deviations (null Q = 0.0751 ± 0.0010; **z = 368**); at combined_score ≥ 700 the separation is larger still (observed Q = 0.6542 over 32 modules against a null of 0.1317 ± 0.0012; **z = 441**). The ≥ 700 re-run recovers 32 modules against the 33 obtained originally — a one-module difference between runs of a stochastic algorithm, which we report rather than reconcile.

A second check is against STRING’s own curated hierarchy. Each data-driven module maps onto a curated cluster carrying the matching functional label, with best-match Jaccard overlap ranging from 0.45 (M0, “Membrane and Signal Transduction”) to 0.83 (M6, “RAC2 GTPase cycle”), and 0.57–0.60 for the ribosome-biogenesis and splicing modules. Global flat agreement is modest in absolute terms (adjusted Rand index 0.18; normalised mutual information 0.27) because STRING’s hierarchy is far finer-grained than our seven major modules (573 candidate clusters); at matched granularity the correspondence is essentially one-to-one. Because those curated clusters are computed from the same underlying associations as our network, we treat this as an internal-consistency check rather than as independent validation; the degree-preserving null above is the test that the partition carries structure beyond the degree sequence.

#### Hubs are constitutive complexes; bottlenecks are nutrient-sensing signalling

The highest-degree nodes are pan-eukaryotic machinery: polyubiquitin-B and -C, 40S/60S ribosomal proteins, RNA polymerase II subunits, glyceraldehyde-3-phosphate dehydrogenase, ATP synthase and proteasome subunits. By contrast, the highest-betweenness nodes — the inter-module information bottlenecks — are overwhelmingly components of the **insulin/IGF → PI3K → AKT → mTOR(RAPTOR) → S6K/FOXO** axis: phosphatidylinositol-4,5-bisphosphate 3-kinase, mTOR, RAC-beta serine/threonine kinase (AKT), ribosomal-protein-S6 kinase, forkhead box protein O (FOXO) and regulatory-associated protein of mTOR (RAPTOR) occupy the top betweenness ranks. Hubs and bottlenecks are largely disjoint — the top 70 by degree and the top 70 by betweenness share only 17 proteins (Figure 1D); hubs sit within stable complexes, bottlenecks bridge between modules — and the distinction sharpens in the physical-only subnetwork (≥ 400: 5,782 proteins, 91,220 edges, modularity 0.61), whose top positions are co-complex members (polyubiquitin, RNA polymerase, ribosomal proteins, the cap-binding complex) while the transient mTOR/AKT regulatory nodes recede.

#### Central classes are consistent with independent functional genetics

Several classes the network ranks most central have been perturbed *in vivo* in work not used in its construction. RNA-interference knockdown of ribosomal-protein genes reduces *Varroa* female offspring and silencing a proteasome subunit affects survival (Huang et al. 2019), so the ribosome-biogenesis (M5) and proteasome (M2) hub classes are experimentally fitness-relevant. A semi-field RNAi screen increased mite infertility by silencing *ptch1*, *ap-1* and *vg1* (Muntaabski et al. 2025); in our network the three *patched homolog 1* paralogues form a coherent signalling cluster and the vitellogenin system appears only through its receptor, consistent with *Varroa*’s partial reliance on host-derived vitellogenin. The reproductive **TOR → vitellogenin** axis, whose mTOR node is a top betweenness bottleneck here, is RNAi-validated in the related mesostigmatid mite *Dermanyssus gallinae*, in which silencing *TOR* or *Vg* reduces fecundity and disrupts embryogenesis (Liu et al. 2024). None of these studies tests an individual predicted edge, and we do not present them as validating the network. What they establish is narrower and still useful: the target classes that orthology transfer alone ranks most central are classes whose perturbation is independently known to impair mite fitness.

#### The network is entirely predicted, and orphan genes are excluded by construction

Two analyses delimit precisely what this network is, and is not.

First, resolving the physical-interaction file into its evidence channels shows that the direct-experimental, database and text-mining scores are uniformly zero throughout, while the transferred channels carry all of the signal. The unthresholded physical subnetwork comprises **203,041 unique undirected edges over 6,905 proteins** (406,082 rows as distributed); at combined_score ≥ 400 it comprises 91,220 edges over 5,782 proteins. The entire *Varroa* interactome is therefore interolog-predicted, with no edge resting on direct experimental support in this organism.

Second, of the 10,241 representative genes, **9,450 carry an eggNOG orthologous group at some level and 791 do not**. Throughout this study “orphan” denotes the absence of an eggNOG orthologous group, not the absence of detectable homology — a distinction we quantify in Results 3. **Not one of the 791 orphans enters the network**, and neither does any gene lacking at least Eukaryota-level conservation: every one of the 7,080 nodes is deeply conserved.

We state plainly that the first half of this observation is definitional rather than empirical. Interolog transfer imports interactions through orthologous relationships, so a gene with no orthologous group can receive no transferred edge; 0 of 791 is the arithmetic of the method, not a discovery about *Varroa*. Three things about it are nevertheless empirical, and they are what the rest of this study measures: the excluded compartment is large and functionally coherent rather than a residue of failed annotation (142 of the 791 orphans carry a substantive product annotation); within the orthology-bearing compartment, where nothing is definitional, visibility does not behave as orthology-transfer intuition predicts; and the same reasoning applied to host comparison produces a quantifiable and asymmetric error rate. An orthology-transfer interactome is a map of what *Varroa* shares with better-studied organisms rather than of what makes it a parasite, and the size and cost of that restriction can be measured.

### 2. A large, distinct compartment of the proteome is invisible to the network

#### Partition by network visibility separates two regimes

Of the 10,241 representative genes, **3,161 (30.9%) lie outside the ≥ 400 network** (Figure 2A). This non-networked fraction is qualitatively unlike the networked one: **62.9% are functionally unannotated** (“uncharacterized”/“hypothetical”) against 11.4% inside the network, and its proteins are markedly shorter (median 330 aa against 478 aa). Short length, poor annotation and absent transferable orthology are together the expected signature of lineage-restricted, fast-evolving genes — but annotation status is itself the dominant correlate of network membership (below), so none of these contrasts should be read as evolutionary before length and annotation are controlled. Within the non-networked fraction, **1,988 genes are unannotated “dark matter”**; this set contains the unannotated orphans and is not disjoint from them (1,988 = 649 unannotated orphans + 1,339 genes that carry an orthologous group but no functional annotation). It is the principal search space for lineage-specific biology.

#### Inside the orthology compartment, visibility is non-monotonic and inverted

Once the definitionally excluded orphans are set aside, the remaining 9,450 orthology-bearing genes allow the question to be put empirically: does network visibility track conservation depth? It does not — and where it varies, it runs against the expected direction. Across conservation levels visibility spans only about 16 percentage points, and the most narrowly resolved bin, genes whose narrowest orthologous group sits at the Acari level, is the **least** visible (74.1% networked), below the Arthropoda-level bin (89.9%), with non-overlapping confidence intervals (Figure 2B). The deficit holds within every length quintile, so it is not a length effect, and it holds within both annotation strata, where it is widest among unannotated genes (35.4% networked at the Acari level against 78.0% at the Arthropoda level).

The reading we take from this is that orthology transfer does not degrade smoothly with evolutionary distance. It behaves as a step function with a hard edge at the presence of an orthologous group, and then imposes a further, counter-intuitive penalty inside the compartment that falls hardest on the most recently resolved groups — precisely the genes sitting closest to the lineage-specific interface. A gradient model, in which visibility falls off gently with decreasing evolutionary depth, would predict the opposite ordering and is not supported.

#### Parasitism-relevant systems are under-represented, with annotation the dominant term

A proteome-wide, keyword-based annotation census of seven functional system classes makes the bias explicit: conserved enzymatic and excitable machinery is almost entirely captured by the network, structural and host-facing systems largely are not. Ranked by the fraction of annotated members lying outside the network, the ordering runs from **cuticle/structural proteins (78.6% outside)** and the **feeding/salivary/secreted set (62.7%)**, through **chitin metabolism (36.4%)** and **neuropeptide ligands (35.0%)**, to **neuropeptide/amine receptors (20.0%)**, **ion channels (10.9%)** and **detoxification enzymes (2.3%)** (Figure 2C).

Two qualifications travel with these figures. First, the ordering is dominated by annotation quality without being reducible to it: in a logistic model of network membership, functional annotation is by far the strongest term (odds ratio 11.24), and the system-class effects survive conditioning on annotation and protein length (cuticle OR 0.077, feeding/salivary 0.099, chitin 0.244; see Materials and Methods). Second, novel salivary and structural genes are unannotated and fall into the dark-matter pool rather than into any class, so these class-level figures **under-state** the lineage-specific load hidden from the network.

#### Host-interface gene families are concentrated outside the network

Multi-copy families relevant to Acari feeding and cuticle biology are heavily non-networked: mucin-5AC (20 paralogues, 75% outside the network), mucin-19 (15), adult-specific rigid cuticular protein 15.7 (14), and cuticle protein 14 (13 paralogues, 92% outside).

We do not read such families as arrays of recent origin. Runs of short, poorly annotated genes are equally the signature of collapsed or mis-split repeats in the assembly, and copy-number counts alone cannot distinguish recent gene birth from assembly artefact. The physical organisation of this compartment is tested against a permutation null in Results 3; the statement supported there is local clustering.

### 3. The lineage-specific compartment is real, secreted, and restricted

#### The orphan set, defined exactly

Of the 10,241 representative genes, **791 carry no eggNOG orthologous group at any taxonomic level**. That is what “orphan” means throughout this study, and the definition is narrower than it sounds: it is a statement about orthologous-group assignment, not about detectable homology. The orphans are an extreme of the non-networked regime — 82% are functionally unannotated and their median length is 226 aa — and three of them are mitochondrially encoded (ATP8, 32 aa; ND4L, 86 aa; ND6, 145 aa), the shortest and most rapidly diverging mitochondrial proteins, whose absence from eggNOG reflects database coverage rather than lineage-specific origin. They are named here because they are excluded from the nuclear-genome analyses below.

An orphan compartment of this size invites three standard objections: that the gene models are annotation errors, that they are assembly artefacts, and that “no orthologous group” has been over-read as “no homologue”. We address them in turn, with evidence that is independent of the orthology assignment that defined the set.

#### The gene models carry the same transcriptional support as networked genes

Mapping all 10,241 proteins back to the genome annotation shows that orphan gene models are supported at the same rate as the rest of the proteome. RNA-seq evidence covers **all annotated introns in 85.2% of orphans, against 91.3% of unannotated non-networked genes, 81.5% of annotated non-networked genes and 89.3% of networked genes**: the qualitative gate is passed at a comparable rate in every compartment (Figure 3A). What differs is the *depth* of that support, which is lower for orphans (median 3 supporting samples against 4 for the networked set) — the expected consequence of lower expression, not of a worse model. Truncated models are rare and do not explain the compartment either (partial=true in 1.3% of orphans against 0.5% of controls).

**Fig. 3.**
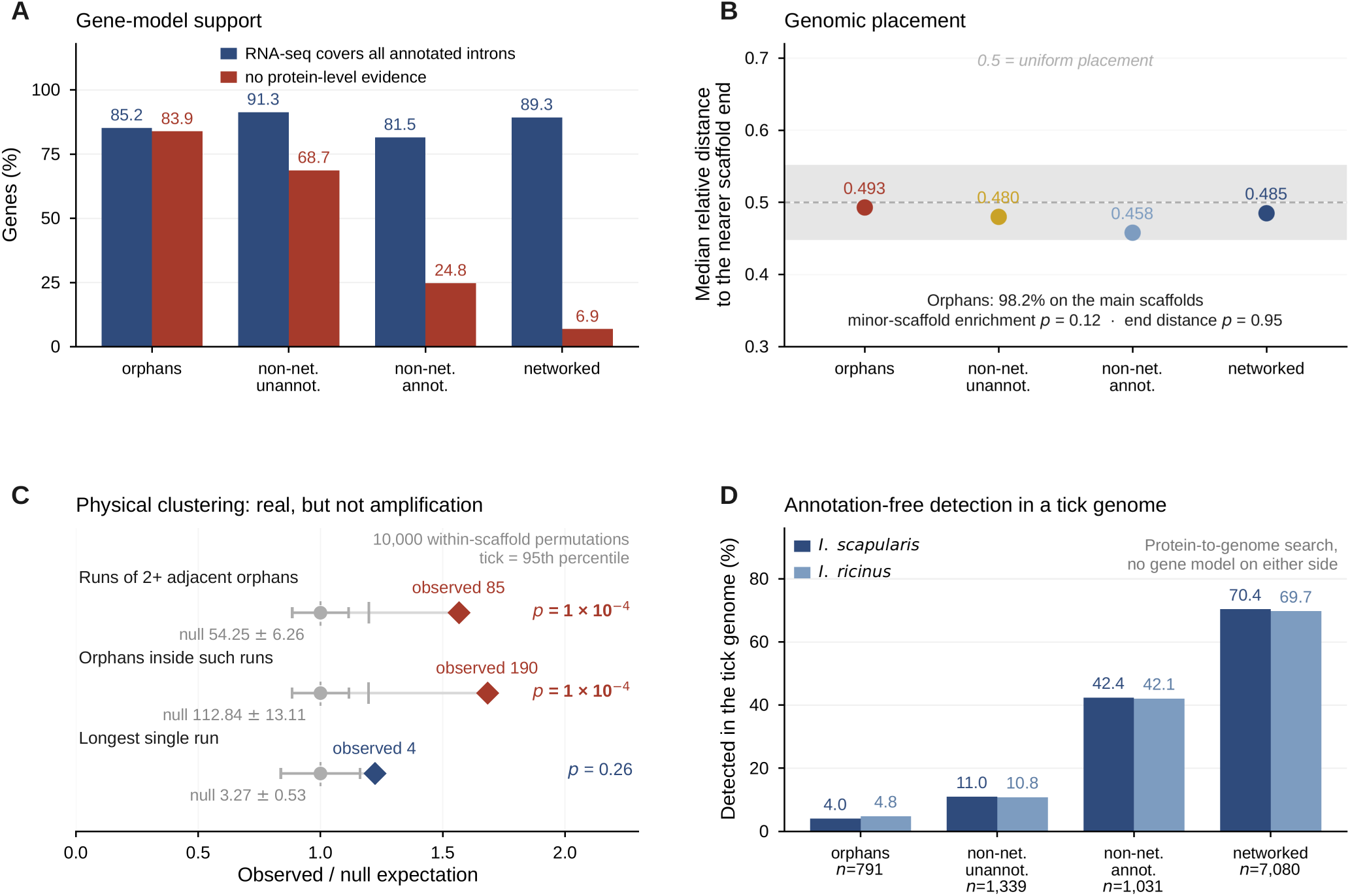
*The excluded compartment is not an artefact of gene prediction, assembly or annotation.* (A) Gene-model support by compartment. RNA-seq covers all annotated introns at a similar rate across all four compartments (orphans 85.2%, unannotated non-networked 91.3%, annotated non-networked 81.5%, networked 89.3%), so orphan gene models are supported no worse than the rest and better than annotated non-networked genes. What differs is protein-level evidence, which is absent for 83.9% of orphans against 6.9% of networked genes. (B) Genomic placement: 98.2% of orphans lie on the main scaffolds with no enrichment on minor ones (Fisher p = 0.12), and their median relative distance to the nearer scaffold end is 0.4929 against 0.4846 for networked genes (Mann–Whitney p = 0.95); 0.5 is the value expected under uniform placement. (C) Physical clustering against 10,000 within-scaffold permutations: 85 runs of two or more adjacent orphans against 54.25 ± 6.26 expected and 190 orphans inside such runs against 112.84 ± 13.11 (both p = 1 × 10⁻⁴), but a longest run of 4 against a null expectation of 3.27 ± 0.53 (p = 0.26) — local clustering, not amplification. (D) Detection by annotation-free protein-to-genome search in two tick genomes, by compartment: in *Ixodes scapularis*, 4.0% of orphans, 11.0% of unannotated non-networked genes, 42.4% of annotated non-networked genes and 70.4% of networked genes; *I. ricinus* gives 4.8%, 10.8%, 42.1% and 69.7%. Because no gene model on either side enters panel D, its gradient cannot be an annotation artefact, and it replicates across two independently assembled genomes. *Alt text:* Four-panel figure. Paired bars comparing four compartments on RNA-seq intron support, which is flat, and on absent protein evidence, which is steep; a point plot of relative distance to scaffold ends clustered around one half with non-significant p-values; a plot of three clustering statistics against their permutation nulls, two significant and one not; and a four-step gradient of genome detectability rising from orphans to networked genes, drawn for two tick genomes.

One contrast that might appear to support the same conclusion should not be used for it. Orphans lack Gnomon protein-similarity support in 83.9% of cases against 8.6% of controls, but protein-similarity support and eggNOG group assignment are both homology signals, and a gene without homologues necessarily lacks both. We report that figure as expected rather than as confirmation; the independent evidence is the RNA-seq column.

#### The orphans are not assembly debris

Scaffold placement excludes the two artefact readings directly. Seven main scaffolds (76.9–32.6 Mb) carry 98.4% of the assembly, with a 215.7-fold drop to the eighth, so the distinction is measured rather than asserted. **98.2% of orphans sit on those main scaffolds** (odds ratio 1.57 for minor scaffolds, **p = 0.12**; 14 genes of 791), which is not what a contamination or low-quality-contig reading predicts. Nor are they concentrated at scaffold margins, where fragmentary models accumulate (**p = 0.95** for relative distance to the nearer scaffold end; Figure 3B). The per-scaffold distribution is deposited; its one departure from uniformity is a *deficit* of orphans, whereas an artefact would predict an enrichment.

Orphans are, however, non-randomly arranged along the scaffolds. Against a within-scaffold permutation null (10,000 permutations), we recover **85 runs of two or more adjacent orphans against an expectation of 54.25 ± 6.26** (p = 1 × 10⁻⁴), with 190 genes in runs against 112.8 expected. The effect is an excess of pairs and triples rather than long arrays: **the longest run is 4 genes against a null expectation of 3.27 (p = 0.26)** (Figure 3C). We therefore describe this as local clustering and not as tandem amplification. The distinction matters, because clustering of short, poorly annotated genes is compatible both with recent gene birth and with collapsed or mis-split repeats in the assembly; this analysis establishes that the clustering exists and does not adjudicate between those readings.

#### “No orthologous group” is not “no homologue”

Domain and secretion evidence recovers structure that the orthology layer does not see. A per-gene domain pass places a recognised domain on **104 of the 791 orphans (13.1%)**. The domains that recur are regulatory rather than parasitism-specific — C2H2 zinc fingers lead with 10 genes — and the parasitism interface appears not as domain annotation but as topology and family structure: **183 orphans carry a signal peptide**, the topology expected of host-facing effectors, and the largest sequence family (17 members, cuticle protein 65-like) is secreted in 15 of 17.

A single-sequence profile search against UniRef50 makes the point more sharply. Some orphans return tens of thousands of hits, and inspection shows why: they carry C2H2 zinc-finger motifs (CxxC…HxxxH), and their hits to octopus, passerines and fish are genuine domain-level homology. A gene can therefore be an orphan in this study’s sense — no orthologous group at any level — while carrying unambiguous homology to distant eukaryotes. We use “orphan” strictly in the orthologous-group sense throughout, and we do not describe any gene in this compartment as newly originated from non-coding sequence: clustering sequences that carry no prior family assignment is not a demonstration of new gene birth, which would require syntenic non-coding ancestry in an outgroup that we have not established.

#### How restricted the orphans are depends on what they are compared against

Two searches of different breadth give different answers, and the difference is the result.

Against the **four comparison proteomes**, 678 of 791 orphans (85.7%) have no detectable homologue, on per-proteome hit counts of *Apis* 0, *Ixodes* 9, *Tropilaelaps* 24 and *Dermanyssus* 91 (Figure 4A). *Tropilaelaps* is phylogenetically closer to *Varroa* than *Dermanyssus* is, so this ordering ranks proteome completeness rather than evolutionary distance — the same panel effect quantified for host-absence calls in Results 4.

**Fig. 4.**
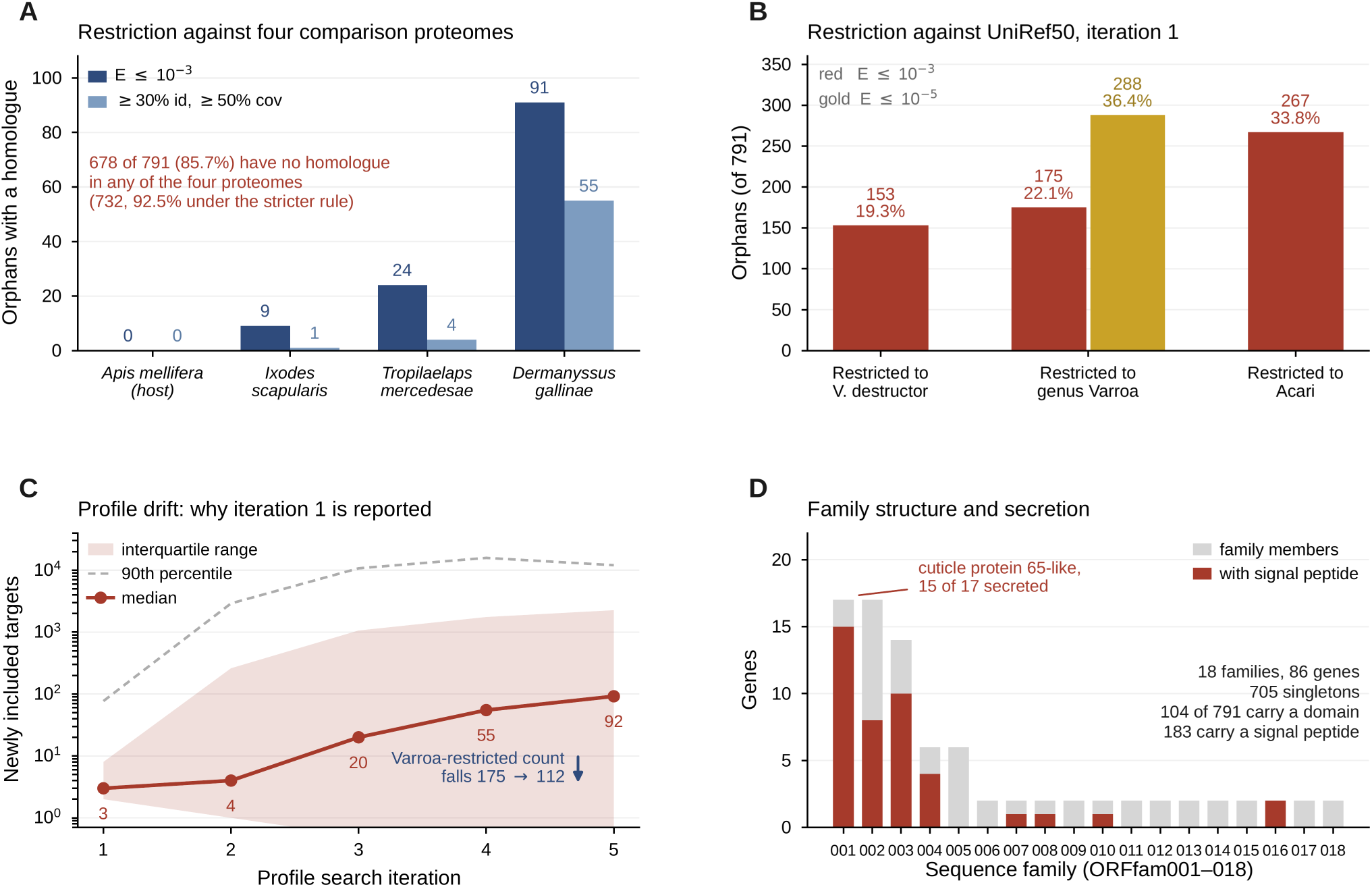
*Within the orphan compartment: restriction depends on what it is measured against.* (A) Restriction of the 791 orphans against four comparison proteomes under two criteria. At E ≤ 1 × 10⁻³ the per-proteome hit counts are *Apis mellifera* 0, *Ixodes scapularis* 9, *Tropilaelaps mercedesae* 24 and *Dermanyssus gallinae* 91, leaving 678 orphans (85.7%) with no detectable homologue in any of the four; under a ≥ 30% identity and ≥ 50% coverage rule the counts fall to 0, 1, 4 and 55, and 732 orphans (92.5%) are restricted. The ordering of the acarine proteomes ranks proteome completeness rather than phylogenetic distance. (B) Restriction against UniRef50 by taxonomic level at iteration 1: 153 orphans (19.3%) with no hit outside *V. destructor*, 175 of 791 (22.1%) outside the genus *Varroa* and 267 (33.8%) outside Acari, with the E ≤ 1 × 10⁻⁵ sensitivity analysis for the genus level shown alongside (288, 36.4%). (C) Why iteration 1 is reported: median newly included targets per round rise from 3 and 4 to 20, 55 and 92, but the ninetieth percentile rises from 77 to over 12,000 — drift is driven by a minority of queries with rapidly growing profiles — and the *Varroa*-restricted count falls from 175 to 112 by iteration 5. (D) Sequence families and secretion: clustering resolves the orphans into 18 families comprising 86 genes plus 705 singletons, the largest being 17 members of a cuticle protein 65-like family of which 15 carry a signal peptide. Across the whole orphan set, 104 of 791 (13.1%) carry a recognised domain and 183 carry a signal peptide. *Alt text:* Four-panel figure. Paired bars of orphan hit counts against four comparison proteomes under two criteria; grouped bars of orphans restricted at three taxonomic levels; a logarithmic plot of newly included search targets per iteration showing a stable median and a sharply rising upper tail; and a bar chart of eighteen sequence families with the secreted fraction of each marked.

Against **UniRef50**, restriction collapses. A single-sequence search (jackhmmer, iteration 1, E ≤ 1 × 10⁻³, release 2026_02, 38,794,121 clusters) places **175 of 791 orphans (22.1%) outside the genus *Varroa***, with 153 (19.3%) outside *V. destructor* and 267 (33.8%) outside Acari (Figure 4B); requiring E ≤ 1 × 10⁻⁵ raises the genus-level figure to 288 (36.4%). The four-proteome estimate over-states genus-level restriction by close to fourfold, and we report this as a correction to our own earlier figure. It carries the same lesson as the host-absence result: absence measured against a small, incompletely annotated sample is absence from the sample.

Iteration depth matters here, and its direction is instructive. Scored at iteration 5 the figure is 112 rather than 175, because profile drift accumulates spurious hits: median newly included targets per round run 3, 4, 20, 55 and 92 while the ninetieth percentile rises from 77 to over 12,000, so drift is driven by a minority of queries with rapidly growing profiles, and only 284 of 790 queries converged within the iteration limit (Figure 4C). Iteration 1 cannot drift and is the level reported throughout: a profile-based search, reasonable to ask for on any orphan claim, fails in its iterated form on exactly this class of short, low-complexity sequence.

#### An annotation-free search confirms the restriction independently

Every comparison above rests on a proteome, and a proteome is an annotation. Searching all representative proteins directly against the *Ixodes* genomes with a protein-to-genome aligner removes that dependency on both sides, and detectability then grades steeply by compartment (Figure 3D):

| compartment | detectable in the *Ixodes* genome |
| --- | --- |
| orphans (nuclear-encoded, n = 788) | **4.0%** |
| non-networked, unannotated | 11.0% |
| non-networked, annotated | 42.4% |
| networked | 70.4% |

Only about one orphan in twenty-five is detectable in a tick genome by a search that uses no annotation at all, against seven in ten networked genes; *I. ricinus* gives the same gradient (4.8%, 10.8%, 42.1% and 69.7%). Because no gene model on either side enters this comparison, the result cannot be an annotation artefact — which is what the objection ultimately requires — and it confirms independently that the orphan compartment is genuinely lineage-restricted.

#### Family structure within the compartment

All-against-all local alignment and community detection resolve the 791 orphans into **18 families comprising 86 genes, with 705 singletons** (Figure 4D). The two largest are secreted and locally clustered: ORFfam001 (17 genes, 14 secreted, annotated “cuticle protein 65-like”) and ORFfam002 (17 genes, 8 secreted, keratin-like), followed by ORFfam003 (14 genes, 8 secreted). The 705 singletons are the paralogue-free core of the compartment.

The secreted, structurally biased signature of the largest orphan families mirrors the cuticle and mucin expansions found among annotated non-networked genes in Results 2, extended into genes with no orthologous group at all. We note two constraints on how far this should be read. The clustering used here is not exactly reproducible (Materials and Methods), and the run lengths reported within families are counted on a different basis from the adjacent-gene runs above, which are the ones with a permutation null attached; where the two disagree, the null-tested figure is the defensible one.

### 4. Host divergence operates in two modes: gene absence and residue-level substitution

A comparative orthology join across the five-species panel places every *Varroa* gene on two axes at once — conservation across Acari, and divergence from the bee. Read on those axes, “divergence from the host” resolves into two processes with almost nothing in common: whole genes the bee lacks, and conserved genes in which a handful of residues differ. The first is where orthology-based methods are used and where they fail, so we quantify that failure before using the compartment it defines.

Two definitions are used throughout, and are stated here because both are permissive. A gene is **Acari-conserved** if it has an orthologous-group counterpart in **at least one** of the three comparison acarine proteomes, not in all three; a gene is a **selectivity candidate** if it is Acari-conserved and called absent from *Apis*. Of 8,747 *Varroa* genes carrying an Acari-level orthologous group, 7,546 (86%) have a counterpart in at least one other acarine.

#### Orthology-identifier absence calls are wrong about two times in three

Declaring a gene absent from the host because its orthologous-group identifier does not appear in the host proteome is the default output of a group-level comparative join. It is also wrong most of the time. Of **2,210 selectivity candidates** called *Apis*-absent on group identity alone, a confirmatory sequence-level search against the *Apis* proteome recovered a sequence homologue for **67.4%** (Figure 5A). The same search against a composition- and length-preserving shuffled null returned no alignment for any of 300 queries (0 of 300; one-sided 95% upper bound 1.0%), so this is not an artefact of a permissive alignment threshold.

**Fig. 5.**
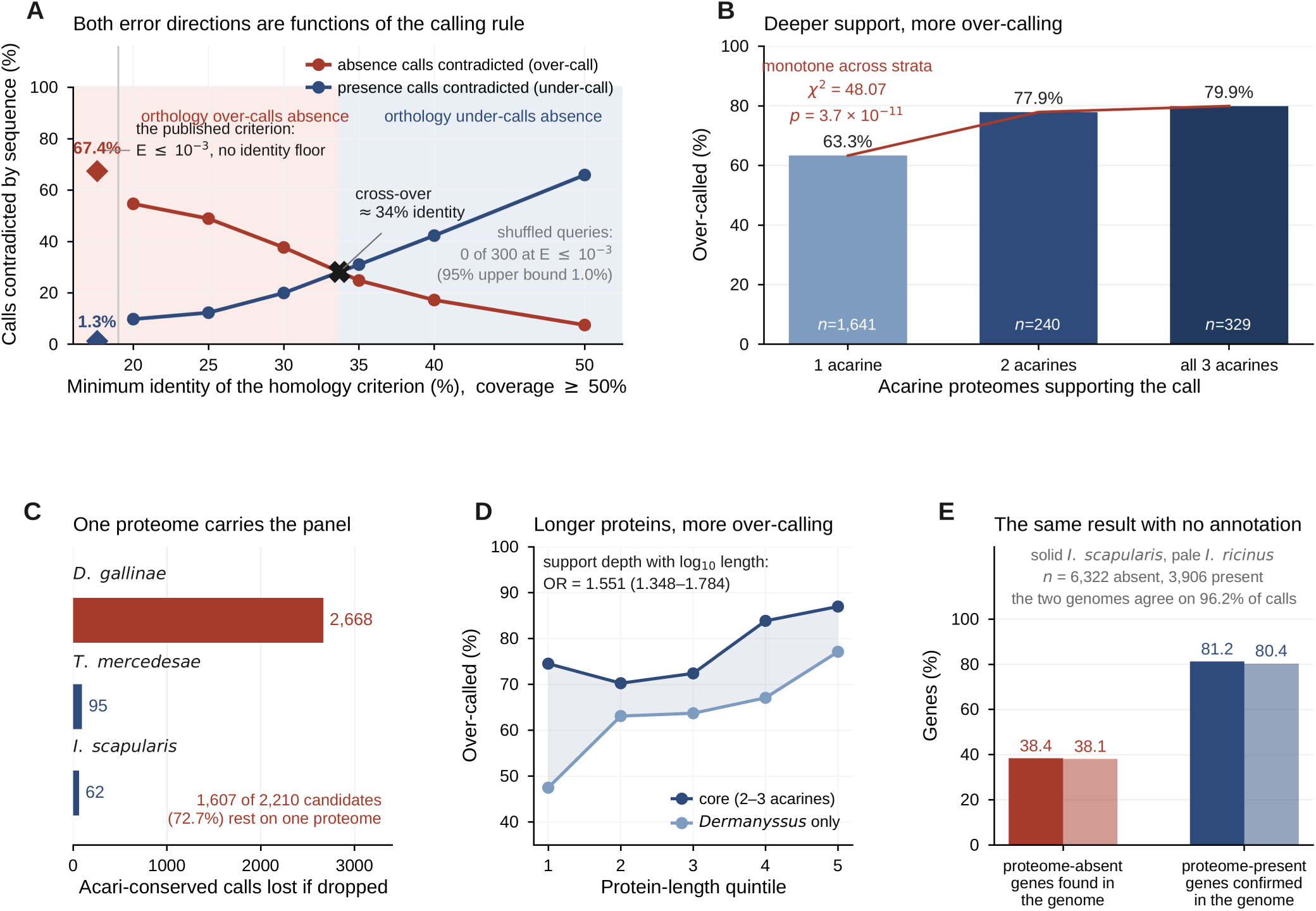
*Orthology-based host-absence calls fail two times in three, and both error directions are functions of the calling rule.* (A) Absence and presence calls contradicted by sequence search, across the homology criterion, on the two populations defined by the same orthology join (2,210 called absent from *Apis*, 5,336 called present). At the published criterion — a DIAMOND alignment at E ≤ 1 × 10⁻³ with no identity or coverage floor, shown as the two diamonds left of the divider — 67.4% of absence calls recover a homologue against 1.3% of presence calls that lose one. Imposing an identity floor moves both, in opposite directions: at ≥ 30% identity over ≥ 50% of the shorter sequence the rates are 37.7% and 20.0%, and at ≥ 50% identity they are 7.5% and 65.9%, so the two curves cross near 34% identity and orthology cannot be said to err in either direction without fixing the criterion first (supplementary table S17; at ≥ 20% identity and ≥ 20% coverage the over-call is 72.4%, so the E-value criterion is not the most permissive available). The E-value criterion is the one carrying a control: none of 300 composition-preserving shuffled queries returned an alignment under identical settings (one-sided 95% upper bound 1.0%), unchanged in DIAMOND’s most sensitive mode. (B) Over-call by acarine support depth, monotone across strata: 63.3% (n = 1,641), 77.9% (n = 240) and 79.9% (n = 329) for calls supported by one, two and three acarine proteomes (χ² = 48.07, p = 3.7 × 10⁻¹¹). (C) Panel composition: leave-one-out removal of *D. gallinae* costs 2,668 Acari-conserved calls against 95 and 62 for the other two acarines, so 1,607 of 2,210 candidates (72.7%) rest on a single proteome. (D) Over-call by protein-length quintile and support stratum. It rises with length in both strata (*Dermanyssus*-only 47.5% to 77.1%; core 74.5% to 87.0%) and the core stratum exceeds the single-proteome stratum in every quintile; acarine support depth retains an odds ratio of 1.551 (95% CI 1.348–1.784) with log₁₀ length in the model. (E) The same asymmetry with no annotation on either side: of 6,322 proteins called absent from the *I. scapularis* proteome, 38.4% align to its genome, while 81.2% of proteome-present genes are confirmed; *I. ricinus* gives 38.1% and 80.4%, and searched identically the two *Ixodes* genomes agree on 96.2% of calls. *Alt text:* Five-panel figure. Two crossing curves showing that the fraction of absence calls contradicted falls, and the fraction of presence calls contradicted rises, as the identity threshold is raised, with the published no-threshold criterion marked separately; a rising three-step plot of over-call against the number of supporting acarine proteomes; a horizontal bar of leave-one-out losses dominated by one species; two rising lines of over-call against length quintile, one stratum above the other throughout; and paired bars showing that proteome-level absence is contradicted by genome-level alignment while proteome-level presence is confirmed, in two tick genomes.

The error is strongly asymmetric: **absence calls fail at 67.4%, presence calls at 1.3%**. Both figures are properties of the calling rule rather than constants. Across 31 criteria computed on the same population (2,210 group-absent and 5,336 group-present Acari-conserved genes), the over-call falls from 67.4% under the E-value rule to **37.7%** at ≥ 30% identity over ≥ 50% coverage and **7.5%** at ≥ 50% identity, while the under-call rises over the same range; the two directions **cross near 34% identity** (28.33% against 28.32% at ≥ 35% identity and ≥ 40% coverage), and at ≥ 20% identity and ≥ 20% coverage the over-call reaches 72.4%, so the E-value rule is not the most permissive criterion available. Orthology cannot be said to err in either direction without fixing the criterion first. Under the matched criterion that carries the shuffled control, orthologous-group identity is serviceable evidence that a gene *is* shared and poor evidence that it is not — a distinction that matters, because the absence direction is the one used to nominate parasite-specific targets.

#### The over-call is not an artefact of panel composition — widening the panel lowered it

The comparison panel is unbalanced and one member dominates it. Proteome coverage runs from *D. gallinae* at 69.96% down through *A. mellifera* (57.47%), *T. mercedesae* (45.87%) and *I. scapularis* (38.14%), and a leave-one-out analysis shows what that costs: dropping *D. gallinae* removes **2,668** Acari-conserved calls, against 95 for *T. mercedesae* and 62 for *I. scapularis* (Figure 5C). Adding *D. gallinae* had grown the candidate set from 603 to 2,210, so **1,607 of 2,210 candidates (72.7%) rest on a single proteome**.

This is a real bias, and it runs opposite to the direction that would flatter the result. Stratified by how many acarine proteomes support a call, the over-call rate is *lowest* in the *Dermanyssus*-only stratum (**63.3%**, n = 1,607) and *highest* in the core supported by *I. scapularis* and/or *T. mercedesae* (**78.3%**, n = 603; Figure 5B), with the pooled figure between them. The rate is monotone in support depth (63.3 → 77.9 → 79.9%; χ² = 48.07, p = 3.7 × 10⁻¹¹) and survives conditioning: with log₁₀ protein length in the model, acarine support depth carries an odds ratio of **1.551** (95% CI 1.348–1.784, p = 8.2 × 10⁻¹⁰), falling only to 1.338 when annotation status is added. The core stratum is higher in **all five** length quintiles without exception.

The practical consequence is the useful part: a narrow panel does not merely add noise to the over-call estimate, it systematically **under**-states it, so the conservative estimate is the 78.3% of the unbiased core rather than the pooled figure. Over-call also rises with protein length, from 47.5% in the shortest quintile to 77.1% in the longest (Figure 5D), with annotation status again the dominant covariate (odds ratio 4.92 here, against 11.24 in the visibility model of Results 2); any single percentage inherits the composition of the set it was drawn from.

#### The same failure appears on the acarine side, in a search that uses no annotation

Both the group-identity call and its sequence-based correction compare proteomes, and a proteome is an annotation. To ask whether absence is a property of the organism or of its gene models, we searched all 10,241 representative proteins directly against both *Ixodes* genomes with a protein-to-genome aligner, using no annotation on either side.

Absence does not survive the test. Of **6,322** proteins called absent from the *I. scapularis* proteome, **2,429 (38.4%) align to the *I. scapularis* genome**; for *I. ricinus* the figure is 2,409 of 6,322 (38.1%). Presence is by contrast stable — 3,172 of 3,906 proteome-present genes are confirmed in the genome (81.2%; 80.4% for *I. ricinus*) — reproducing on the acarine side the same asymmetry found on the host side (Figure 5E).

The two tick genomes also supply the within-genus control that the panel composition invites. Searched identically, *I. scapularis* and *I. ricinus* agree on **96.2%** of calls (Jaccard 0.933; 5,381 shared of 5,601 and 5,548), so substituting one *Ixodes* species for the other cannot have driven any result reported here; the Materials and Methods record why the substitution was unavoidable.

Within this annotation-free frame, detectability is steeply graded by compartment membership, from 4.0% of orphans to 70.4% of networked genes. That gradient is the independent evidence that the orphan compartment is genuinely lineage-restricted, and it is reported with the orphan controls in Results 3.

#### After correction, a secreted and Acari-conserved compartment remains

Requiring **both** group-level and sequence-level absence from *Apis* leaves **1,715 genes** that are robustly Acari-conserved and host-absent (1,513 from the orthology join, 202 recovered by the sequence layer; Figure 6A).

**Fig. 6.**
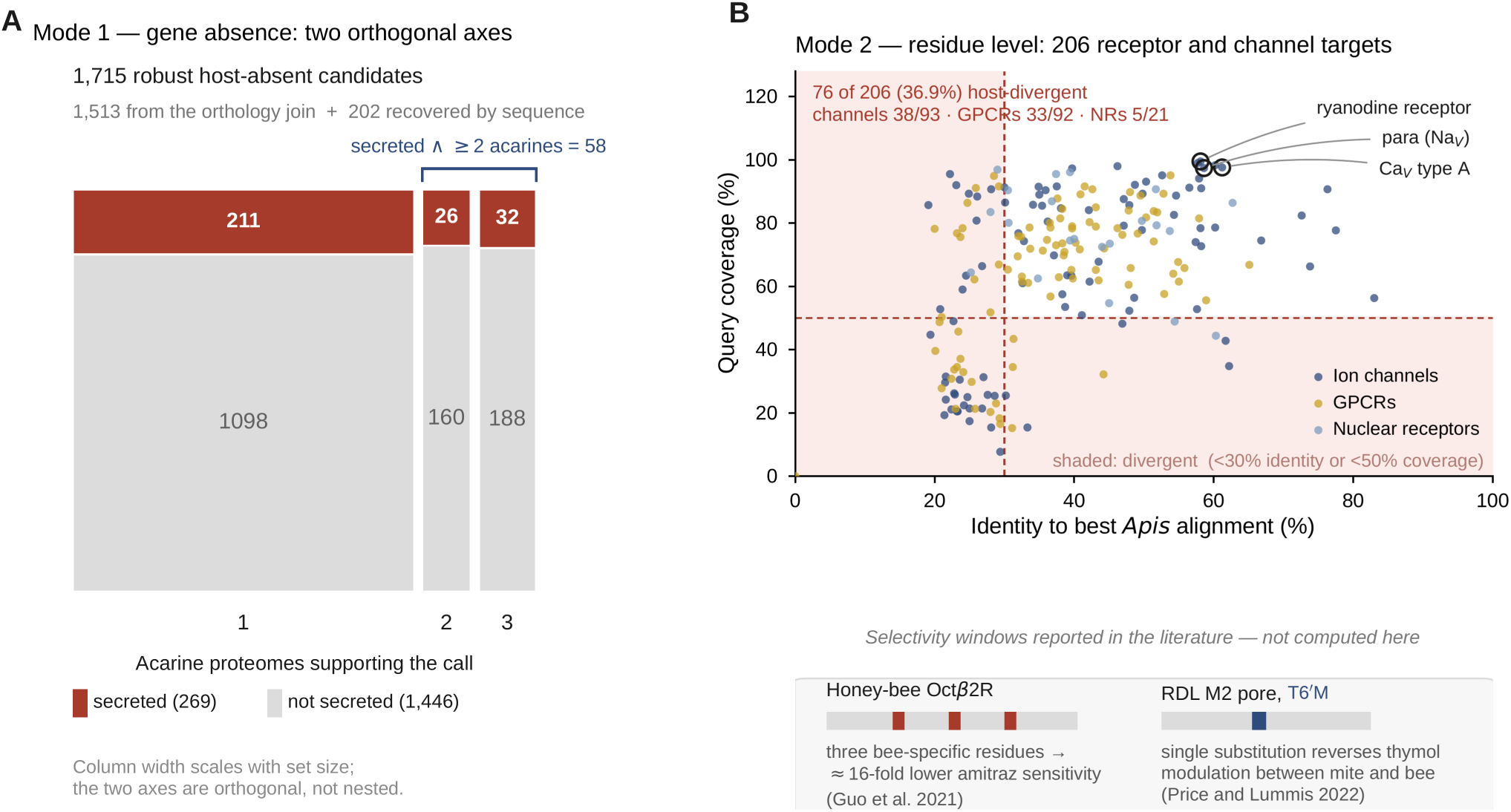
*Host divergence operates in two modes, and the two select for different properties.* (A) Mode 1, gene absence. Requiring both orthologous-group absence and sequence absence from *Apis* leaves 1,715 robust candidates (1,513 from the orthology join, 202 recovered by the sequence layer). These resolve on two orthogonal axes, not a funnel: 269 candidates are secreted and 220 are conserved in all three acarine proteomes, and the two sets intersect in only 32 genes; a third intersection, secreted candidates conserved in at least two acarine proteomes, contains 58. The secreted fraction is near-constant across support depth (16.1%, 14.0% and 14.5% for one, two and three acarines), so the two axes are close to independent rather than nested. (B) Mode 2, residue level. All 206 receptor and channel targets plotted by identity and coverage of their best *Apis* alignment; 76 (36.9%) are host-divergent, defined as below 30% identity or below 50% query coverage — 38 of 93 ion channels, 33 of 92 G-protein-coupled receptors and 5 of 21 nuclear receptors. The three canonical acaricide targets are among the most conserved proteins in the repertoire: the voltage-gated sodium channel *para* (58.6% identity over 97.4% of its length), the ryanodine receptor (58.1%, 99.5%) and the voltage-gated calcium channel (61.2%, 97.6%). The insets show the three bee-specific residues that render the honey-bee Octβ2R approximately 16-fold less amitraz-sensitive (Guo et al. 2021) and the single M2-pore substitution T6′M that reverses thymol modulation of RDL between mite and bee (Price and Lummis 2022). Both are taken from the cited literature and were not computed here. *Alt text:* Two-panel figure. A mosaic of 1,715 host-absent candidates split by secretion and by the number of supporting acarine proteomes, with the 58-gene and 32-gene intersections marked; and a scatter of 206 receptor and channel targets by identity and coverage against the bee, with the divergent region shaded and three named acaricide targets circled in the conserved corner, beside schematics of a three-residue receptor window and a single-residue pore substitution.

These resolve on two axes that are largely independent, not on a nested series of filters. **269 of the candidates are secreted** and **220 are conserved in all three acarine proteomes**, but the two sets **intersect in only 32 genes**. A third intersection, secreted candidates conserved in at least two acarine proteomes, contains 58. The axes select for different properties, and presenting them as successive narrowings of a single funnel would misrepresent both. The leading families are the expected host-facing effector classes — insect cuticle RR proteins (PF00379), CBM14 chitin-binding mucins, invertebrate-type lysozyme, and, most cleanly, a **dermonecrotic phospholipase-D/SicTox-like family present in the tick and all three mites yet absent from the bee**.

The orphan compartment shows the same panel effect from the other direction (Results 3): narrow panels over-call lineage specificity for the same reason they under-state host-absence over-call.

#### Mode 2 — for the conserved target channels, selectivity is residue-level

The classic acaricide-target channels behave in the opposite way. The voltage-gated sodium channel (*para*), the ryanodine/IP₃ receptor and the voltage-gatedcalcium channel return **zero** whole-protein host-divergent hits: all are pan-arthropod-conserved and present in the bee. Across the receptor and channel repertoire only 76 of 206 genes (36.9%) are sequence-confirmed host-divergent at the whole-protein level (Figure 6B), and these fall in the most divergent GPCR and ionotropic-glutamate families rather than at the validated target sites.

For those sites, selectivity is not a property of gene presence but of individual residues, and the published pharmacology pins it to the residue. Two examples, both taken from that literature rather than computed here, are given with their numbers in Figure 6: three bee-specific residues make the honey-bee Octβ2R roughly **16-fold** less sensitive to amitraz than its *Varroa* counterpart (Guo et al. 2021), and a single M2-pore substitution, **T6′M**, reverses the direction in which thymol modulates RDL between mite and bee (Price and Lummis 2014, 2022).

In both cases the protein is conserved and present in both species, yet a two-or three-residue difference opens a real selectivity window. The two modes are therefore not two descriptions of one process: Mode 1 is visible to orthology-based methods and is precisely where they make their characteristic error, while Mode 2 is invisible to them altogether, because at the level of gene presence there is nothing to see.

## Discussion

We set out to measure what an orthology-transfer interactome omits in a non-model parasite, and what the same orthology logic costs when it is used to compare that parasite with its host. The omission is not a gap but a second, evolutionarily distinct compartment of the proteome; the comparative cost is a large, asymmetric and panel-dependent error rate. Together they organise the *Varroa* proteome into two compartments and host divergence into two processes, with consequences for how comparative interactomes are read, for the evolutionary biology of parasite proteomes, and — secondarily — for the control of an economically critical pest.

### Network centrality and lineage-specific parasitism are near-orthogonal axes

The definitional part belongs first, because it is what a sceptical reader should press on. That none of the 791 orphans enters the network is arithmetic rather than discovery: interolog transfer imports interactions through orthologous relationships, so a gene with no orthologous group can receive no transferred edge, and we claim nothing from 0 of 791 beyond the boundary it draws.

Everything on either side of that boundary is empirical. The excluded compartment is large (3,161 genes, 30.9% of the proteome), coherent, and not a residue of failed annotation: 142 orphans carry a substantive product annotation; their gene models carry RNA-seq support across all annotated introns at a rate comparable to every other compartment (85.2% against 89.3% for networked genes); they sit on the main scaffolds and not at scaffold margins; and a protein-to-genome search using no annotation on either side — and therefore immune to the annotation objection — recovers them in a tick genome at 4.0% against 70.4% for networked genes. Inside the orthology-bearing compartment, where nothing is definitional, visibility then fails to behave as orthology-transfer intuition predicts. It runs in the wrong direction, the most narrowly resolved bin (Acari, 74.1% networked) sitting below the Arthropoda bin (89.9%) with non-overlapping intervals and the deficit holding within every length quintile. Orthology transfer behaves as a step function with a hard edge at the presence of an orthologous group, and then penalises the genes nearest that edge — precisely those closest to the lineage-specific interface. The consequence for study design is the finding rather than a corollary of it: centrality is rich for an ancient conserved core and empty for a fast-evolving secreted interface, so neither map alone recovers the organism.

### Host divergence is two evolutionary processes, not one

The comparative join sharpens this into an evolutionary statement. Divergence from the host decomposes into gene absence and residue-level substitution, and the two are concentrated in different compartments. The gene-absence mode builds an Acari-conserved, secreted compartment — cuticle RR proteins, CBM14 chitin-binding mucins, invertebrate-type lysozyme, and most cleanly a dermonecrotic phospholipase-D/SicTox-like family present in the tick and all three mites yet absent from the bee — together with a genuinely *Varroa*-restricted orphan core. The residue-level mode operates on the opposite class of genes: the canonical target channels (*para*, the ryanodine/IP₃ receptor, the voltage-gated calcium channel) show no whole-protein host divergence at all, and their selectivity lives instead in two- and three-residue windows, as the amitraz/Octβ2R and thymol/RDL pharmacology demonstrates at single-residue resolution (Guo et al. 2021; Price and Lummis 2022). That a parasite should diverge from its host both by retaining secreted effectors the host never had and by tuning shared machinery at a handful of positions is a coherent two-process picture, and one that any single whole-protein method would collapse into one misleading signal. A spider-venom peptide supplies an instructive coda: Ht1a is mite-selective on topical application while inhibiting the *Varroa* and *Apis* sodium channels with comparable potency (Herzig et al. 2026), a reminder that residue-level conservation does not exhaust the routes to differential vulnerability.

### A general caution for non-model comparative genomics

Two results here generalise beyond *Varroa*, and both concern absence.

The first is that an orthology-transfer interactome omits the lineage-specific compartment of any non-model organism *silently*: the missing genes never appear as missing, nothing in the output marks the boundary, and the omission is largest exactly where the interest is greatest — in parasites, symbionts and rapidly evolving lineages.

The second is measurable, and larger than we expected. Of 2,210 genes called absent from the host on orthologous-group identity alone, 67.4% recover a sequence homologue in the host proteome. The same search against composition-preserving shuffled sequences recovers none, so the rate is a property of the calling rule and not of a permissive alignment threshold. The error is strongly asymmetric — 67.4% of absence calls fail against 1.3% of presence calls — and it falls in the worst place, because the absence direction is the one used to nominate host-specific genes, gene losses and selective targets.

Three further analyses say how such a number should be reported. First, it depends on the panel, and not gently. Ours is dominated by one proteome, and the bias runs against the result rather than for it: over-call is *lowest* in the *Dermanyssus*-only stratum (63.3%) and *highest* in the core supported by the other two acarines (78.3%), monotone in support depth and robust to conditioning on protein length and annotation status. The direction is unsurprising once stated — a gene present in more acarine proteomes is more likely genuinely conserved, hence more likely to have a host homologue after all — but its consequence is not: widening a panel does not add noise to an over-call estimate, it systematically *understates* it.

Second, the rate depends on composition. Over-call rises from 47.5% in the shortest length quintile to 77.1% in the longest, and annotation status is the dominant covariate here as in the model of network visibility. Any single percentage inherits the composition of the candidate set it was drawn from, and should travel with it rather than as a constant.

Third, the same failure appears where no annotation is used on either side: of 6,322 proteins called absent from the *Ixodes scapularis* proteome, 38.4% align to the *I. scapularis* genome, while 81.2% of proteome-present genes are confirmed — the host-side asymmetry reproduced on the acarine side, with gene models removed from the comparison altogether.

The transferable form of the lesson is that both panel-dependent quantities in this study moved by a large factor in the direction the panel predicts, and in opposite directions. Restriction measured against four proteomes placed 678 of 791 orphans (85.7%) beyond detectable homology; measured against UniRef50 the genus-restricted figure is 175 of 791 (22.1%). Narrow panels over-call lineage specificity and under-state host-absence over-call for the same reason: absence measured against a small and incompletely annotated sample is absence from the sample, not from the clade. Two rules follow, and neither is expensive. Treat group-level identity as evidence that a gene is shared and not that it is absent, confirming every absence call at the sequence level; and report any absence-derived statistic with the panel it was measured against, because the panel, not the threshold, is the term that moves it.

One methodological caveat belongs here rather than in a footnote. A reader may reasonably ask why the orphan set was not characterised with an iterated profile search; as Results 3 shows, the iterated form drifts on exactly this class of short, low-complexity sequence. Every restriction level reported here is therefore scored at iteration 1, a plain single-sequence search that cannot drift, and its recovery of homologues of the three mitochondrially encoded orphans, which must have them, is the built-in control that it is configured correctly.

### Implications for control (broader impact)

The architecture described here maps onto target discovery and explains why the two halves of the proteome need different methods. The conserved end offers RNAi-tractable targets — the mTOR/FOXO bottleneck, the ribosome-biogenesis hubs and the TOR → vitellogenin axis — essential, but shared with the bee and so requiring residue-level selectivity engineering. The lineage-specific end offers the bee-sparing targets: the salivary effectors, of which the chitinase Vd-CHIsal is required for feeding and survival (Becchimanzi et al. 2020), with further candidates since identified from the salivary-gland transcriptome (Becchimanzi et al. 2025); the secreted phospholipase-D/SicTox, cuticle, mucin and lysozyme families; and the host-absent neuropeptide systems. Selectivity is achievable at either end but free at neither: gene-absent targets are selective by construction and uncharacterised, conserved targets well understood but in need of residue-by-residue tuning. Delivery constrains both, since environmental RNAi in mites can be localised rather than systemic (Bensoussan et al. 2022).

The acaricide-resistance literature falls along the same axis, and where it falls is itself informative. Metabolic resistance runs through the conserved detoxification families: reduced cytochrome-P450 proinsecticide activation underlies coumaphos resistance (Vlogiannitis et al. 2021b), a specific P450 is associated with amitraz and flumethrin resistance (Mavridis et al. 2025), and RNAi knockdown of an ABC-transporter efflux pump raises amitraz mortality (Ricigliano et al. 2026). Target-site resistance runs through the conserved channels and receptors: the sodium channel *para*, where pyrethroid mutations are widespread in treated populations (Vlogiannitis et al. 2021a; Celikkol and Dogac 2025; Yalcin et al. 2026), the octopamine receptor and the RDL channel. Both arms sit inside the compartment that orthology transfer sees perfectly well, which is the point rather than an aside: half a century of chemical control has selected on the part of the proteome comparative methods map most reliably, while the bee-sparing opportunities lie in the compartment those methods cannot see. The one conspicuous lineage-level signal in the enzyme repertoire is an acetylcholinesterase expansion to 14 genes (supplementary table S10), the shared acarine reservoir for organophosphate and carbamate chemistry. Only 9 of the 14 are in the network, so even this expansion is partly invisible to the methods mapping the conserved compartment. The amitraz axis is the one system in which mechanism, resistance genetics and a selectivity window all coincide.

Two cautions belong with any use of this map. Individual resistance substitutions are not interchangeable with selection signals, and the catalogue assembled here is a literature synthesis at family level with no selection test. We propose no new molecule or protocol; the contribution is the map and the evolutionary reading of it.

### Limitations

Six caveats bound the interpretation. (1) Every interaction is interolog-predicted; no edge rests on direct experimental support in this organism, so individual edges are hypotheses. We make no claim about the shape of the degree distribution, which this study does not test and which most empirical networks fail to satisfy under formal criteria (Broido and Clauset 2019). (2) *Apis*-absence is reliable only after sequence confirmation, and we applied that correction to the candidate sets rather than exhaustively to every raw group-absence call. (3) Domain and secretion calls are precomputed topology heuristics, not experimental localisation, so “secreted candidate” remains a prediction. (4) The panel is unbalanced: 72.7% of candidates rest on the *D. gallinae* proteome alone, and although the stratified analysis shows this dilutes rather than inflates the over-call, absolute *Dermanyssus*-only conservation counts remain the least secure figures here. (5) The tick tier is *I. scapularis* rather than *I. ricinus*, which has no annotated proteome; searched identically against both genomes the two species agree on 96.2% of calls (Jaccard 0.933), so the substitution cannot have driven any result reported here. (6) The restriction figures are search-dependent by construction — 22.1% genus-restricted at E ≤ 1 × 10⁻³, 36.4% at E ≤ 1 × 10⁻⁵, 85.7% against the four-proteome panel — and we carry the conservative one.

Two further constraints are matters of provenance rather than inference and are stated in Materials and Methods: the keyword patterns behind the system-class census and the seed behind the orphan-family clustering were not preserved, so neither is exactly reproducible, although both outputs are deposited.

## Conclusion

The *Varroa destructor* proteome is organised into two evolutionarily distinct compartments — an ancient, conserved, network-embedded core, and a secreted, lineage-restricted interface that orthology transfer cannot see — and it diverges from its honey-bee host by two mechanisms, gene absence and residue-level substitution, that fall in those two compartments respectively. Reconstructing one map yields a systematically incomplete picture; trusting orthology-based host-absence calls without sequence confirmation yields a partly false one, at a rate measured here at 67.4% and 78.3% in the least biased stratum. The two-axis, two-mode view provides both a scaffold of conserved targets and a sequence-defined catalogue of lineage-specific, host-facing candidates, and a transferable caution for the comparative genomics of any non-model organism whose biology lives in its fastest-evolving genes.

## Materials and Methods

### Genome, proteome and identifier backbone

All analyses are anchored on the RefSeq assembly Vdes_3.0 of *Varroa destructor* (GCF_002443255.2; Techer et al. 2019). To obtain one protein per protein-coding gene, the RefSeq proteome was collapsed to **10,241 representative sequences** by selecting, for each gene, the longest isoform with a deterministic tie-break. A CDS protein_id ↔ GeneID map derived from the genome GFF served as the identifier backbone for every downstream join. Unless stated otherwise, “gene” denotes one of these 10,241 representatives.

### Interactome reconstruction

Because *V. destructor* is not a core STRING organism and no experimental protein–protein interaction data exist for it, a network was obtained by submitting the representative proteome to STRING v12.0 (Szklarczyk et al. 2023), which builds an entirely prediction-based, interolog-transfer network from a chosen reference clade — here Acari (taxon 6933). The submission corresponds to STRING organism identifier **STRG0A76ODC** (pipeline 12.0.2). The STRING flat files analysed, including the physical-interaction file with its direct/transferred evidence-channel split retained, are listed in the deposit manifest.

STRING node identifiers were de-prefixed to RefSeq accessions. STRING distributes each interaction as two directed rows; these were collapsed, and all edge counts reported here are unique undirected pairs. Networks were built at two fixed confidence thresholds — combined_score ≥ 400 (medium confidence, primary) and ≥ 700 (high confidence) — in python-igraph v1.0.0 (Csárdi and Nepusz 2006). Edge weight was defined as combined_score/1000 and path distance as 1 − weight. For every node we computed degree, weighted strength, weighted betweenness, closeness and eigenvector centrality. Communities were detected with the Leiden algorithm under the modularity objective (Traag et al. 2019) using leidenalg v0.12.0 with a fixed random seed. Per-module functional over-representation (KEGG, Reactome, Gene Ontology biological process and molecular function) was assessed by the hypergeometric test against the network background with Benjamini– Hochberg control of the false-discovery rate; per-term adjusted values are deposited.

### Module validation: curated clusters and a degree-preserving null

Two independent checks were applied to the Leiden partition.

First, each module was compared with STRING’s own precomputed hierarchical clusters, and best-match overlap was quantified by the Jaccard index. Because those curated clusters are derived from the same underlying associations as the network itself, this is an internal-consistency check and is reported as one.

Second, and independently of STRING, modularity was tested against a **degree-preserving null**: 100 randomised graphs generated by edge rewiring with 10 swap attempts per edge, preserving the degree sequence exactly, with Leiden run identically on each. The observed modularity is reported as a z-score against that null distribution at both confidence thresholds. Because Leiden is stochastic, module counts can differ by one small module between runs; the re-run count is reported alongside the original where they differ.

### Conservation layer and the definition of “orphan”

For each gene we recorded the narrowest eggNOG (Huerta-Cepas et al. 2019) taxonomic level at which an orthologous group is defined, along the lineage root → Eukaryota → Opisthokonta → Metazoa → Bilateria → Arthropoda → Arachnida → Acari, and retained the Acari-level (6933) and Arthropoda-level (6656) group identifiers for cross-species joining. Genes with no eggNOG orthologous group at any level were defined as **orphans** (n = 791).

This definition is narrower than the word implies and is used consistently throughout: an orphan is a gene with no orthologous-group assignment, **not** a gene with no detectable homology. Of the 791, 104 carry a recognised protein domain, and some return large numbers of database hits through shared domain content. No gene is described in this study as newly originated from non-coding sequence, which would require syntenic non-coding ancestry in an outgroup that we have not established.

### Domain and secretion annotation

Each representative was annotated against UniProtKB (taxon 109461; UniProt Consortium 2023) parsed for InterPro (Paysan-Lafosse et al. 2023) and Pfam (Mistry et al. 2021) entries, signal-peptide predictions (SignalP; Teufel et al. 2022) and transmembrane-helix predictions (Phobius; Käll et al. 2004), together with RefSeq and GeneID cross-references. Genes were matched first by direct RefSeq accession and otherwise by a gene-level fallback via NCBI GeneID (same-length isoform preferred), with the match route recorded as a provenance column. A gene was scored a **secreted candidate** when a signal peptide was present and no transmembrane helix was predicted. Where a count of signal-peptide-carrying genes is reported without that second condition, it is stated as such. These topology calls are precomputed heuristics, not experimental localisation, and are reported as such throughout.

### Gene-model support and genomic placement of the orphan set

Two analyses were run directly on the genome annotation, independently of any homology evidence.

#### Gene-model support

All 10,241 representative proteins were mapped to the genomic GFF (protein_id → parent RNA → mRNA attributes). For each model we recorded whether RNA-seq evidence in the model_evidence attribute covers all annotated introns, the number of supporting samples, and the partial flag.

The four compartments were compared on the qualitative gate and, by Mann– Whitney U, on support depth. The inference attribute does not occur on mRNA features in this annotation, so the analysis rests on model_evidence alone.

Gnomon protein-similarity support was recorded but is **not** used as evidence, because protein-similarity support and eggNOG group assignment are both homology signals and a gene without homologues necessarily lacks both.

#### Genomic placement

Scaffold lengths were measured from GFF region features. Seven scaffolds carrying 98.4% of the assembly were designated main scaffolds, the cut being set by a measured 215.7-fold drop in length to the eighth. We tested enrichment of orphans on minor scaffolds (Fisher’s exact test), relative distance to the nearer scaffold end (Mann–Whitney U), per-scaffold distribution (χ²) and physical clustering against a within-scaffold permutation null: runs of two or more adjacent orphans separated by ≤ 100 kb, counted against 10,000 permutations of orphan status within each scaffold preserving gene positions and per-scaffold counts (seed 42), testing both the number of runs and the longest run. Three mitochondrially encoded orphans (NP_758876.1/ATP8, NP_758882.1/ND4L, NP_758883.1/ND6) were identified and are excluded from nuclear-genome analyses.

### Sequence-level restriction of the orphan set

Restriction was assessed at two levels of breadth, because the two give materially different answers.

#### Four-proteome panel

The 791 orphans were searched with DIAMOND (Buchfink et al. 2021) against the *Apis mellifera*, *Ixodes scapularis*, *Tropilaelaps mercedesae* and *Dermanyssus gallinae* proteomes at E ≤ 1 × 10⁻³. This threshold was validated by the shuffled-sequence control described under “Panel composition and the over-call benchmark”.

#### UniRef50

The same 791 sequences were searched against UniRef50 (Suzek et al. 2015) release 2026_02 (38,794,121 clusters, md5 3228886e9d749f050f60e9a0ce1f727d) with jackhmmer from HMMER v3.4 (Eddy 2011) at E ≤ 1 × 10⁻³, and hits were mapped to taxonomy with the NCBI taxdump of 2026-08-05. **All reported restriction levels are scored at iteration 1**, which is a plain single-sequence search and cannot drift. Iterated scoring was evaluated and rejected: the median number of newly included targets per round rises sharply from the third iteration, only a minority of queries converge within the iteration limit, and profile drift moves genes out of the restricted class by accumulating spurious hits. A sensitivity analysis at E ≤ 1 × 10⁻⁵ is reported alongside the primary threshold. Of the 791 queries, 790 returned a hit table; one (XP_022656634.1) did not, and is reported separately rather than counted as having no homologue.

Scoring refuses to proceed unless every search table terminates with a completion marker. This guard was added after an earlier run in which memory-killed jobs were logged as complete; a truncated table is indistinguishable from a homologue-free query and would have inflated the restricted count precisely for the queries with the most hits.

### Annotation-free protein-to-genome search

To test absence without relying on annotation on either side, all 10,241 representative proteins were aligned directly to the *I. scapularis* (GCF_016920785.2; De et al. 2023) and *I. ricinus* (GCA_964198075.1) genome assemblies with miniprot v0.18 (Li 2023), requiring ≥ 30% identity over ≥ 50% coverage; the full threshold grid is deposited. Genes were then cross-tabulated by proteome-level call (present/absent) against genome-level detection, and separately by network-visibility compartment. Agreement between the two *Ixodes* genomes, searched identically, provides a within-genus control on the panel substitution. Of the 791 orphans, 788 are nuclear-encoded and entered this search; the three mitochondrially encoded orphans were excluded, because alignments of mitochondrial proteins to nuclear scaffolds represent nuclear mitochondrial insertions rather than orthologues.

*I. scapularis* was used in place of the *I. ricinus* originally specified. This was a necessity rather than a preference: as checked on 2026-08-05, *I. ricinus* has nine assemblies, all contig-or scaffold-level and none annotated, so no proteome exists. The within-genus control above quantifies the consequence directly.

### Sequence clustering of the orphan set

The 791 orphan sequences were subjected to all-against-all local alignment with parasail (Daily 2016) using sw_trace_striped_16 (BLOSUM62; gap open 11, gap extend 1), restricted to sequence pairs within a 2.5× length band. An edge was drawn when percent identity ≥ 0.35, coverage of the shorter sequence ≥ 0.60 and aligned length ≥ 40 aa, with edge weight = identity × coverage.

Families were defined as Louvain communities (Blondel et al. 2008) at resolution 1.0. Run lengths reported within these families are counted on a different basis from the adjacent-gene runs of the placement analysis above; where the two disagree, the permutation-tested figure is the one used.

### Comparative orthology join and the host-divergence axis

The comparative panel comprised five species: **(A)** *V. destructor* (query); **(B)** *I. scapularis* (taxon 6945; GCF_016920785.2); **(C)** *D. gallinae*, whose predicted proteome (Burgess et al. 2018) was submitted to STRING as build **STRG0A97ZCG** to obtain an eggNOG-namespace orthology file; **(C′)** *T. mercedesae* (taxon 418985; Dong et al. 2017); and **(D)** *A. mellifera* (host; taxon 7460; Amel_HAv3.1 / GCF_003254395.2; Wallberg et al. 2019), the selectivity reference. Inputs were the STRING v12.0 protein.orthology and protein.sequences files for taxa 7460, 6945 and 418985 and for submission STRG0A97ZCG, plus the per-gene annotated *Varroa* backbone and the representative FASTA.

Two complementary layers covered all 10,241 representatives. **Layer 1** joined genes on shared eggNOG orthologous-group identifiers at the Acari (6933) and Arthropoda (6656) levels; presence and group copy number were recorded for each comparison species. **Layer 2** sequence-searched the 1,041 genes lacking a group at either level (791 orphans plus 250 root-only genes) against each species proteome with parasail sw_stats_striped_16 (BLOSUM62; gap 11/1), using a prefilter of ≥ 3 shared distinct 5-mers and a target length within 0.5–2.0× of the query; a hit required ≥ 30% identity over ≥ 50% coverage **of the shorter sequence**.

#### Definitions

A gene is **Acari-conserved** if it is present in at least one of the three comparison acarine proteomes — not in all three. A gene is a **selectivity candidate** if it is Acari-conserved and called absent from *Apis*. Both definitions are permissive and are stated because they materially affect every downstream count; both were recovered exactly against the deposited join matrices.

#### The *Apis*-absence rule

Because orthologous-group-identifier matching systematically over-calls absence from *Apis* (quantified below), every host-divergence and selectivity call required **both** group-absence **and** sequence-absence in *Apis*.

### Panel composition and the over-call benchmark

Two analyses quantify how the comparison panel itself affects the results.

#### Leave-one-out

Each acarine proteome was removed in turn and acari_conserved and the selectivity-candidate set recounted, giving the marginal contribution of each proteome. Proteome coverage — the fraction of that species’ proteome carrying an eggNOG group at the relevant level — was computed for all four comparison species.

#### Over-call benchmark

Selectivity candidates called *Apis*-absent by group identity alone were re-searched against the *Apis* proteome with DIAMOND at E ≤ 1 × 10⁻³, and the proportion recovering a sequence homologue is reported as the over-call rate. *Shuffled-sequence control.* To establish that the E-value criterion does not recover alignments from composition alone, 300 of the 2,210 group-absent queries were shuffled with amino-acid composition and length preserved, then searched against the same *Apis mellifera* database under identical settings (DIAMOND v2.2.4, E ≤ 1 × 10⁻³). No query returned an alignment (0 of 300; one-sided 95% upper bound 1.0%), unchanged under --very-sensitive. The 300 shuffled sequences are deposited verbatim (A3_null_shuffled.faa, md5 80b482a91467667a304794fad0dff2e8) together with the search output, so the control is reproducible exactly without re-running the shuffle; the seed itself was not preserved. Candidates were stratified by the number of supporting acarine proteomes, and the over-call rate tested for a trend across strata by χ². Because both length and annotation status could confound this, a logistic model of over-call was fitted with acarine support depth, log₁₀ protein length and annotation status as terms, and the stratum contrast was additionally examined within length quintiles. The presence direction was benchmarked in the same way, so that the two error rates are directly comparable. The over-call rate depends on the homology criterion, and the dependence was quantified rather than assumed: both error directions were recomputed from the same search output over a grid of 31 identity-and-coverage criteria (deposited as supplementary table S17). The E-value rule is the criterion carrying the shuffled control and is the one reported as the headline; the grid is reported alongside it because the two directions cross within the range of criteria in ordinary use.

### Target-family assignment

Representative genes were assigned to functional subsystems and families by a priority-ordered rule set combining Pfam domain identity with RefSeq product keywords. The first matching rule won, so each gene received at most one family assignment. The complete rule set, with the Pfam accession behind every family, is deposited.

### Literature integration

Functional and resistance evidence was layered from a curated literature set with all DOIs verified against publisher records. Functional support is recorded at family level — a single *Varroa* resistance study lifts all members of that family equally — and is reported as such rather than as gene-resolved validation.

### Figures and reproducibility

Figures were generated with matplotlib (Hunter 2007). Resolution is encoded in each figure filename; gene labels are shown in full without truncation. All analytical steps are scripted, and the node, edge, module, enrichment, comparative-join, orphan and target tables are provided as machine-readable files (see Data availability).

### Reproducibility limitations

Three limitations on exact reproducibility are stated here rather than left to be discovered.

First, the analyses were run in two phases separated in time, and the earlier computing environment was not preserved. Where a figure could not be reproduced exactly, that is stated at the point of use; environment details for both phases are in the deposit README.

Second, the keyword patterns underlying the system-class annotation census were lost with the earlier environment. The census is reported as originally computed, from the deposited summary table. The class assignment used as a covariate in the logistic model of network visibility is a **separately derived** classification and is not the same object as the census table; both are deposited so that they can be compared directly.

Third, the community-detection step that defines the orphan families used a random seed that was not recorded, so the exact partition is not reproducible. The families, their sizes and the resulting family/singleton split are deposited as computed.

Raw intermediate outputs of the profile search are not deposited because of their size; they are regenerable from the stated release, md5 and search parameters.

### Use of AI assistance

A large language model was used to assist with language editing of the manuscript and to support data-processing and analysis scripting. All output was reviewed, verified, and approved by the author, who takes full responsibility for the manuscript’s content.

## Supplementary material

Supplementary material is available at *Genome Biology and Evolution* online: supplementary tables S1–S17 and their index, uploaded as separate files.

## Data availability

All data underlying this study are publicly available. The analysis scripts, the network node and edge tables with per-channel evidence scores, the module and module-validation tables, the degree-preserving null distributions, the orphan annotation, gene-model-support, genomic placement and sequence-clustering tables, both comparative-join matrices, the leave-one-out and over-call benchmark tables, the protein-to-genome search results and the target-family assignments are deposited at Zenodo under DOI 10.5281/zenodo.21933932, together with a VERSION.txt recording the release and checksum of every database used. Versioned code is additionally available at https://github.com/stepanryba-prog/varroa-gbe/.

The *Varroa destructor* protein-interaction network was obtained by submitting the representative proteome to STRING v12.0 under organism identifier STRG0A76ODC (pipeline 12.0.2), and the *Dermanyssus gallinae* proteome under STRG0A97ZCG. Because user submissions to STRING are not guaranteed to persist, the complete node, edge and evidence-channel tables returned by both submissions are included in the Zenodo deposit, so that every analysis reported here can be reproduced without re-submission.

Genome and proteome accessions: *Varroa destructor* GCF_002443255.2 (Vdes_3.0); *Ixodes scapularis* GCF_016920785.2; *Ixodes ricinus* GCA_964198075.1; *Apis mellifera* GCF_003254395.2 (Amel_HAv3.1); *Tropilaelaps mercedesae* NCBI taxon 418985. Comparative orthology and sequence files were obtained from STRING v12.0; UniRef50 release 2026_02 (md5 3228886e9d749f050f60e9a0ce1f727d) and the NCBI taxonomy dump of 2026-08-05 were used as distributed. Raw intermediate outputs of the profile search are not deposited because of their size; they are regenerable from the stated release, checksum and search parameters. No new sequence data were generated by this study.

## Acknowledgements

None.

## Author contributions

Štěpán Ryba is the sole author. He conceived and designed the study, performed all analyses, and wrote the manuscript.

