## Supplemental tables for "Orthology transfer maps only the conserved core of the *Varroa destructor* proteome and over-calls host absence two times in three"

Štěpán Ryba

Tables S2, S3, S5, S10, S13, S15, S16 and S17 are reproduced in full below. The remaining tables are per-gene matrices of 175 to 10,241 rows and up to 62 columns, which cannot be rendered legibly on a printed page; they are deposited in machine-readable form at Zenodo (DOI 10.5281/zenodo.21933932) and are listed in the index with their row and column counts. Column dictionaries for every table are given in the deposited index file.

### Index of supplementary tables

| Table | Title | Data rows | Columns | In this PDF |
|---|---|---|---|---|
| S1 | Annotated interactome nodes (combined_score >= 400) | 7080 | 27 | Zenodo only |
| S2 | System-by-system visibility census | 7 | 5 | yes |
| S3 | Non-networked-enriched gene-family expansions (>=5 paralogues) | 63 | 5 | yes |
| S4 | Annotated orphan genes (no eggNOG OG at any level) | 791 | 24 | Zenodo only |
| S5 | Orphan sequence families | 18 | 11 | yes |
| S6 | Comparative orthologous-group join (Layer 1, all 10, 241 genes) | 10241 | 22 | Zenodo only |
| S7 | Comparative sequence search (Layer 2, OG-unjoinable genes) | 1041 | 26 | Zenodo only |
| S8 | Ranked robust selectivity candidates | 1715 | 16 | Zenodo only |
| S9 | Enzyme / metabolism target repertoire (Phase 1) | 175 | 34 | Zenodo only |
| S10 | Enzyme / metabolism family summary | 21 | 11 | yes |
| S11 | Phase-1 Apis sequence-homology confirmation | 175 | 7 | Zenodo only |
| S12 | Receptor / channel / signalling target repertoire (Phase 2) | 206 | 62 | Zenodo only |
| S13 | Receptor / channel family summary | 22 | 6 | yes |
| S14 | Phase-2 Apis sequence-homology confirmation | 206 | 6 | Zenodo only |
| S15 | Phase-1 curated literature evidence | 63 | 8 | yes |
| S16 | Phase-2 curated literature evidence | 23 | 7 | yes |
| S17 | Bidirectional over-call and under-call across 31 homology criteria | 31 | 10 | yes |

*All 17 tables are also provided as tab-separated files in the Zenodo deposit.*

### Table S2. System-by-system visibility census

Annotated genes per functional system class, split by network membership. *pct_invisible* is the percentage of annotated members of the class lying outside the interactome.

| system_class | annotated_total | in_network | outside_network | pct_invisible |
|---|---|---|---|---|
| Chitin metabolism | 11 | 7 | 4 | 36.4 |
| Cuticle/structural | 70 | 15 | 55 | 78.6 |
| Feeding/salivary/secreted | 67 | 25 | 42 | 62.7 |
| Neuropeptide ligands | 20 | 13 | 7 | 35 |
| Neuropeptide/amine receptors | 15 | 12 | 3 | 20 |
| Ion channels | 46 | 41 | 5 | 10.9 |
| Detoxification(GST/P450/ABC) | 43 | 42 | 1 | 2.3 |

### Table S3. Non-networked-enriched gene-family expansions (≥ 5 paralogues)

Product-name families with at least five paralogues, ranked by the fraction lying outside the network.

| fam | copies | in_network | outside_network | pct_outside |
|---|---|---|---|---|
| mucin-5ac | 20 | 5 | 15 | 75 |
| nose resistant to fluoxetine protein 6 | 18 | 8 | 10 | 55.6 |
| mucin-19 | 15 | 7 | 8 | 53.3 |
| zinc finger protein 512b | 15 | 3 | 12 | 80 |
| adult-specific rigid cuticular protein 15.7 | 14 | 5 | 9 | 64.3 |
| cuticle protein 14 | 13 | 1 | 12 | 92.3 |
| histone h2b | 11 | 0 | 11 | 100 |
| histone h2a | 11 | 0 | 11 | 100 |
| histone h4 | 10 | 10 | 0 | 0 |
| toll receptor tollo | 10 | 10 | 0 | 0 |
| organic cation transporter protein | 10 | 4 | 6 | 60 |
| calphotin | 10 | 1 | 9 | 90 |
| down syndrome cell adhesion molecule protein dscam2 | 10 | 6 | 4 | 40 |
| alpha-tocopherol transfer protein | 10 | 1 | 9 | 90 |
| elongation of very long chain fatty acids protein aael008004 | 9 | 3 | 6 | 66.7 |
| mfs-type transporter slc18b1 | 9 | 0 | 9 | 100 |
| flocculation protein flo11 | 9 | 2 | 7 | 77.8 |
| glucose dehydrogenase [fad%2c quinone] | 8 | 6 | 2 | 25 |
| glutamate receptor ionotropic%2c kainate 2 | 8 | 7 | 1 | 12.5 |
| glutamate-gated chloride channel | 8 | 8 | 0 | 0 |
| acetylcholinesterase-1 | 8 | 6 | 2 | 25 |
| adult-specific rigid cuticular protein 15.5 | 8 | 4 | 4 | 50 |
| histone-lysine n-methyltransferase smyd3 | 8 | 8 | 0 | 0 |
| elongation of very long chain fatty acids protein 4 | 7 | 1 | 6 | 85.7 |
| hemicentin-2 | 7 | 6 | 1 | 14.3 |
| elongation of very long chain fatty acids protein 7 | 7 | 3 | 4 | 57.1 |
| cathepsin l1 | 7 | 5 | 2 | 28.6 |
| long-chain-fatty-acid--coa ligase 4 | 7 | 7 | 0 | 0 |
| innexin inx2 | 7 | 6 | 1 | 14.3 |
| af4/fmr2 family member 4 | 7 | 2 | 5 | 71.4 |
| glycine-rich cell wall structural protein | 6 | 0 | 6 | 100 |
| sodium-coupled monocarboxylate transporter 1 | 6 | 0 | 6 | 100 |
| trypsin-1 | 6 | 5 | 1 | 16.7 |
| muscle lim protein 1 | 6 | 0 | 6 | 100 |
| peroxidase | 6 | 4 | 2 | 33.3 |
| sialin | 6 | 6 | 0 | 0 |
| leukocyte surface antigen cd53 | 6 | 6 | 0 | 0 |
| histone h3 | 6 | 6 | 0 | 0 |
| monoacylglycerol lipase abhd12 | 6 | 0 | 6 | 100 |
| nephrin | 6 | 6 | 0 | 0 |
| carbonic anhydrase-related protein 10 | 6 | 0 | 6 | 100 |
| cuticle protein 65 | 6 | 0 | 6 | 100 |
| insulin-degrading enzyme | 6 | 5 | 1 | 16.7 |
| acid-sensing ion channel 1 | 5 | 3 | 2 | 40 |
| collagen alpha-1(i) chain | 5 | 2 | 3 | 60 |

| fam | copies | in_network | outside_network | pct_outside |
|---|---|---|---|---|
| cuticle protein 38 | 5 | 0 | 5 | 100 |
| cuticle protein 63 | 5 | 0 | 5 | 100 |
| gamma-aminobutyric acid receptor subunit beta | 5 | 5 | 0 | 0 |
| extensin | 5 | 1 | 4 | 80 |
| dentin sialophosphoprotein | 5 | 2 | 3 | 60 |
| adam 17 protease | 5 | 5 | 0 | 0 |
| camp-dependent protein kinase catalytic subunit alpha | 5 | 4 | 1 | 20 |
| multidrug resistance-associated protein 1 | 5 | 4 | 1 | 20 |
| nardilysin | 5 | 4 | 1 | 20 |
| scavenger receptor class b member 1 | 5 | 5 | 0 | 0 |
| sodium-dependent glucose transporter 1a | 5 | 0 | 5 | 100 |
| serine/arginine repetitive matrix protein 1 | 5 | 2 | 3 | 60 |
| proline-rich extensin protein epr1 | 5 | 0 | 5 | 100 |
| sphingomyelin phosphodiesterase | 5 | 5 | 0 | 0 |
| transcription factor spt20 homolog | 5 | 2 | 3 | 60 |
| stearoyl-coa desaturase 5 | 5 | 5 | 0 | 0 |
| uncharacterized transmembrane protein ddb_g0289901 | 5 | 2 | 3 | 60 |
| zinc finger protein 865 | 5 | 0 | 5 | 100 |

### Table S5. Orphan sequence families

Sequence families recovered by clustering the 791 orphan proteins. *max_tandem_run* is the longest run of family members adjacent on one scaffold; *n_SP* counts predicted signal peptides and *n_secreted* signal peptide without transmembrane helix. The community-detection seed was not recorded (see Materials and Methods, Reproducibility limitations).

| family | size | n_scaffolds | max_ tandem_run | n_SP | n_secreted | len_ min | len_ max | pfam | example | members |
|---|---|---|---|---|---|---|---|---|---|---|
| ORFfam001 | 17 | 6 | 5 | 15 | 14 | 74 | 545 | | cuticle protein 65-like | XP_022658946.1, XP_022662934.1, XP_022652222.1, XP_022659826.1, XP_022670726.1, XP_022647712.1, XP_022654595.1, XP_022661622.1, XP_022657854.1, XP_022661727.1, XP_022662923.1, XP_022659128.1, XP_022659152.1, XP_022657856.1, XP_022658894.1, XP_022643595.1, XP_022660099.1 |
| ORFfam002 | 17 | 5 | 2 | 8 | 8 | 77 | 248 | | keratin, type I cytoskeletal 9-like | XP_022647989.1, XP_022672824.1, XP_022663463.1, XP_022673540.1, XP_022646626.1, XP_022662845.1, XP_022651743.1, XP_022660857.1, XP_022653721.1, XP_022647682.1, XP_022643335.1, XP_022650655.1, XP_022658252.1, XP_022645174.1, XP_022648170.1, XP_022664432.1, XP_022648611.1 |
| ORFfam003 | 14 | 6 | 1 | 10 | 8 | 80 | 411 | | uncharacterized protein F12A10.7-like | XP_022645744.1, XP_022658138.1, XP_022643499.1, XP_022647044.1, XP_022670562.1, XP_022647755.1, XP_022648570.1, XP_022669416.1, XP_022647530.1, XP_022653064.1, XP_022652134.1, XP_022649584.1, XP_022656632.1, XP_022666843.1 |
| ORFfam004 | 6 | 3 | 2 | 4 | 4 | 174 | 480 | | proline-rich protein 4-like | XP_022649129.1, XP_022643973.1, XP_022652920.1, XP_022662080.1, XP_022655527.1, XP_022651887.1 |
| ORFfam005 | 6 | 4 | 1 | 0 | 0 | 89 | 307 | | putative uncharacterized protein DDB_G0294196 | XP_022663458.1, XP_022645302.1, XP_022644261.1, XP_022667771.1, XP_022656634.1, XP_022653617.1 |
| ORFfam006 | 2 | 2 | 0 | 0 | 0 | 110 | 137 | | uncharacterized protein LOC111243073 | XP_022643859.1,XP_022672469.1 |
| ORFfam007 | 2 | 2 | 0 | 1 | 1 | 197 | 237 | | uncharacterized protein LOC111245980 | XP_022650828.1,XP_022644001.1 |
| ORFfam008 | 2 | 2 | 0 | 1 | 1 | 269 | 315 | | histidine-rich glycoprotein-like | XP_022644016.1,XP_022658189.1 |
| ORFfam009 | 2 | 1 | 2 | 0 | 0 | 312 | 548 | | uncharacterized protein LOC111245095 isoform X1 | XP_022648664.1,XP_022647676.1 |

| family | size | n_scaffolds | max_ tandem_run | n_SP | n_secreted | len_ min | len_ max | pfam | example | members |
|---|---|---|---|---|---|---|---|---|---|---|
| ORFfam010 | 2 | 1 | 2 | 1 | 1 | 344 | 614 | | collagen alpha-2(IV) chain-like | XP_022651770.1,XP_022651771.1 |
| ORFfam011 | 2 | 2 | 0 | 0 | 0 | 273 | 305 | | uncharacterized protein LOC111252599 isoform X1 | XP_022666508.1,XP_022653157.1 |
| ORFfam012 | 2 | 1 | 2 | 0 | 0 | 296 | 349 | PF16087 | uncharacterized protein LOC111247784 isoform X1 | XP_022654868.1,XP_022654867.1 |
| ORFfam013 | 2 | 1 | 2 | 0 | 0 | 411 | 411 | | uncharacterized protein LOC111249036 | XP_022658056.1,XP_022660774.1 |
| ORFfam014 | 2 | 1 | 2 | 0 | 0 | 348 | 393 | | uncharacterized protein LOC111249437 | XP_022659040.1,XP_022659089.1 |
| ORFfam015 | 2 | 2 | 0 | 0 | 0 | 103 | 125 | | uncharacterized protein LOC111251194 | XP_022663294.1,XP_022649692.1 |
| ORFfam016 | 2 | 1 | 2 | 2 | 2 | 150 | 150 | | uncharacterized protein LOC111251496 | XP_022663863.1,XP_022663290.1 |
| ORFfam017 | 2 | 2 | 0 | 0 | 0 | 185 | 185 | | uncharacterized protein LOC111255058 isoform X1 | XP_022672420.1,XP_022658822.1 |
| ORFfam018 | 2 | 1 | 2 | 0 | 0 | 288 | 288 | | uncharacterized protein LOC111243543 | XP_022645022.1,XP_022673385.1 |

### Table S10. Enzyme / metabolism family summary

*n_seq_divergent* counts genes below the identity and coverage threshold against *Apis mellifera*; *n_OGabsent_unconf* counts orthologous-group absence calls not confirmed by sequence search.

| subsystem | family | n_ genes | n_ networked | n_secreted | n_seq_ divergent | n_ OGabsent_ unconf | varroa_ genes | max_ score | top_gene | top_product |
|---|---|---|---|---|---|---|---|---|---|---|
| Detoxification | GST | 15 | 15 | 0 | 5 | 6 | 15 | 0.764 | XP_022651584.1 | glutathione S-transferase kappa 1-like |
| Detoxification | CCE_esterase | 8 | 8 | 4 | 5 | 2 | 8 | 0.731 | XP_022655373.1 | isoamyl acetate-hydrolyzing esterase 1 homolog |
| Detoxification | ABC_transporter | 41 | 25 | 0 | 7 | 6 | 41 | 0.698 | XP_022661600.1 | ATP-binding cassette sub-family F member 2-like |
| Detoxification | AKR | 3 | 3 | 0 | 2 | 0 | 3 | 0.676 | XP_022649228.1 | uncharacterized protein LOC111245286 |
| Detoxification | Aldehyde_DH | 13 | 13 | 0 | 1 | 2 | 13 | 0.666 | XP_022664694.1 | aldehyde dehydrogenase family 16 member A1-like isoform X1 |
| Antioxidant | GPx | 5 | 5 | 1 | 0 | 3 | 5 | 0.664 | XP_022669930.1 | epididymal secretory glutathione peroxidase-like |
| Detoxification | CYP_P450 | 30 | 29 | 1 | 6 | 3 | 30 | 0.649 | XP_022649681.1 | uncharacterized protein LOC111245503 isoform X1 |
| Antioxidant | Thioredoxin | 8 | 7 | 0 | 0 | 1 | 8 | 0.646 | XP_022660754.1 | thioredoxin-1-like |
| Antioxidant | SOD | 5 | 5 | 1 | 0 | 1 | 5 | 0.644 | XP_022653269.1 | superoxide dismutase [Mn], mitochondrial-like |
| Neurotransmitter | MAO | 2 | 2 | 0 | 2 | 2 | 2 | 0.641 | XP_022668486.1 | amine oxidase [flavin-containing]-like isoform X1 |
| Antioxidant | Catalase | 1 | 1 | 0 | 0 | 0 | 1 | 0.63 | XP_022661620.1 | catalase-like isoform X1 |
| Neurotransmitter | GABA_T | 1 | 1 | 0 | 0 | 0 | 1 | 0.609 | XP_022667072.1 | 4-aminobutyrate aminotransferase, mitochondrial-like isoform X1 |
| Antioxidant | Peroxiredoxin | 5 | 4 | 1 | 1 | 1 | 5 | 0.595 | XP_022659002.1 | peroxiredoxin-6-like |
| Detoxification | Epoxide_hydrolase | 1 | 1 | 0 | 0 | 0 | 1 | 0.568 | XP_022668682.1 | epoxide hydrolase 1-like |
| Neurotransmitter | AChE | 14 | 9 | 6 | 1 | 10 | 14 | 0.549 | XP_022654432.1 | acetylcholinesterase-1-like |
| Detoxification | UGT | 2 | 2 | 0 | 0 | 1 | 2 | 0.541 | XP_022656245.1 | galactosylgalactosylxylosylprotein 3-beta-glucuronosyltransferase 3-like |
| Neurotransmitter | DBH_TBH | 1 | 1 | 1 | 0 | 0 | 1 | 0.54 | XP_022668942.1 | dopamine beta-hydroxylase-like isoform X1 |
| Neurotransmitter | ChAT | 1 | 1 | 0 | 0 | 0 | 1 | 0.518 | XP_022650492.1 | choline O-acetyltransferase-like |
| Detoxification | Sulfotransferase_cyt | 14 | 11 | 0 | 3 | 6 | 14 | 0.511 | XP_022668161.1 | heparan sulfate 2-O-sulfotransferase 1-like |
| Neurotransmitter | DDC_decarb | 3 | 3 | 0 | 0 | 1 | 3 | 0.486 | XP_022648094.1 | aromatic-L-amino-acid decarboxylase-like |

| subsystem | family | n_ genes | n_ networked | n_secreted | n_seq_ divergent | n_ OGabsent_ unconf | varroa_ genes | max_ score | top_gene | top_product |
|---|---|---|---|---|---|---|---|---|---|---|
| Detoxification | FMO | 2 | 2 | 0 | 0 | 0 | 2 | 0.475 | XP_022653664.1 | dimethylaniline monooxygenase [N-oxide-forming] 5-like |

### Table S13. Receptor / channel family summary

*n_host_divergent* counts genes whose best *Apis* alignment falls in the divergent region of the identity–coverage plane (Fig. 6B).

| subsystem | family | n_genes | n_networked | n_secreted | n_host_divergent |
|---|---|---|---|---|---|
| GPCRs | GPCR_neuropeptide | 33 | 26 | 0 | 9 |
| Ion channels | iGluR_NMDA_kainate | 25 | 15 | 0 | 15 |
| GPCRs | GPCR_secretin_adhesion | 23 | 18 | 0 | 13 |
| Nuclear receptors | NR_other | 17 | 16 | 0 | 4 |
| GPCRs | GPCR_aminergic | 15 | 13 | 0 | 3 |
| GPCRs | GPCR_rhodopsin_other | 10 | 8 | 0 | 4 |
| Ion channels | LGIC_nAChR | 10 | 8 | 0 | 5 |
| Ion channels | LGIC_GABA_RDL | 9 | 9 | 0 | 1 |
| GPCRs | GPCR_glutamate_GABAB | 9 | 9 | 0 | 2 |
| Ion channels | Kv_K | 9 | 8 | 0 | 1 |
| Ion channels | LGIC_GluCl | 9 | 8 | 0 | 4 |
| Ion channels | VGIC_other | 7 | 5 | 0 | 5 |
| Ion channels | LGIC_glycine_5HT3 | 5 | 5 | 0 | 1 |
| Nuclear receptors | NR_EcR_RXR_USP | 4 | 4 | 0 | 1 |
| Ion channels | TRP | 4 | 4 | 0 | 2 |
| Ion channels | Cl_channel_other | 4 | 4 | 0 | 3 |
| Ion channels | VGCC_Ca | 3 | 3 | 0 | 0 |
| Ion channels | LGIC_other_cysloop | 2 | 1 | 0 | 1 |
| Ion channels | CNG_HCN | 2 | 2 | 0 | 0 |
| Ion channels | RyR_IP3R | 2 | 2 | 0 | 0 |
| Ion channels | VGSC_Na | 2 | 2 | 0 | 0 |
| GPCRs | GPCR_opsin | 2 | 2 | 0 | 2 |

### Table S15. Phase-1 curated literature evidence

Curated evidence supporting the enzyme and metabolism target families. Bibliographic strings were harvested automatically and are inconsistent between records; the DOI is authoritative and the harvested *journal* and *title* fields are therefore not reproduced here. They are retained verbatim in the deposited file.

| doi | year | first_ author | families | organisms | evidence |
|---|---|---|---|---|---|
| 10.1007/s11356-019-05247-2 | 2019 | Sabová | GST;Detox_general | Varroa;Apis(host) | biochemical/functional;toxicology |
| 10.1016/j.pestbp.2020.104567 | 2020 | Glavan⁎ | AChE;GST;CCE_esterase;Octopamine_R | Varroa;Apis(host) | biochemical/functional |
| 10.1016/j.compbiolchem.2011.03.004 | 2011 | — | GST;CYP_P450;CCE_esterase | Ixodes | descriptive |
| 10.1038/s41598-020-79057-9 | 2020 | Genath | CYP_P450;Detox_general | Varroa;Apis(host) | transcriptomic |
| 10.1101/415356 | 2018 | Rinkevich | CYP_P450;CCE_esterase | Varroa;Apis(host) | biochemical/functional |
| 10.1186/s13071-026-07461-7 | 2026 | Ricigliano | ABC_transporter;Octopamine_R | Varroa;Apis(host) | RNAi/knockdown;resistance/target-site;transcriptomic |
| 10.3390/biology9090237 | 2020 | Morfin | AChE;CCE_esterase | Varroa;Apis(host) | descriptive |
| 10.1016/j.pestbp.2024.106156 | 2024 | Li | AChE;GST;CYP_P450;ABC_transporter;CCE_esterase;Detox_general | Varroa;Apis(host) | transcriptomic |

| doi | year | first_ author | families | organisms | evidence |
|---|---|---|---|---|---|
| 10.1038/s41598-025-06466-z | 2025 | Fahrbach | Octopamine_R | Varroa;Apis(host) | descriptive |
| 10.1016/j.pestbp.2025.106682 | 2025 | Liu | AChE;GST;CYP_P450;ABC_transporter;CCE_esterase | Dermanyssus | RNAi/knockdown;biochemical/functional |
| 10.1016/j.envres.2021.111836 | 2022 | Wu | GST;CYP_P450;CCE_esterase;Detox_general | Varroa;Apis(host) | toxicology |
| 10.1016/j.actatropica.2020.105763 | 2021 | Hernandez | GST | Ixodes;other-tick | descriptive |
| 10.1186/s13071-015-0960-9 | 2015 | Bartley | AChE;GST;CYP_P450;CCE_esterase | Dermanyssus;Ixodes;other-tick | descriptive |
| 10.1002/ps.8434 | 2024 | Hernández-Rodríguez | Octopamine_R | Varroa;other-tick;Apis(host) | resistance/target-site |
| 10.1016/j.vetpar.2024.110121 | 2024 | Wang | GST;CYP_P450;ABC_transporter;Detox_general | Dermanyssus | RNAi/knockdown |
| 10.1016/j.ijpara.2017.03.007 | 2017 | Barrero | AChE;CYP_P450;CCE_esterase | Ixodes;other-tick | descriptive |
| 10.1016/j.vetpar.2023.110028 | 2023 | Alimi | AChE;CCE_esterase | Dermanyssus | descriptive |
| 10.3390/vaccines12020148 | 2024 | Win | GST | Varroa;Dermanyssus;other-tick | biochemical/functional |
|  | 2005 | Rudenko | GST | Ixodes;other-tick | transcriptomic |
| 10.1016/j.pestbp.2020.104603 | 2020 | Vu | Cl_channel | Varroa | descriptive |
| 10.1016/j.pestbp.2025.106537 | 2025 | — | AChE;CYP_P450;CCE_esterase;Octopamine_R;Cl_channel | Varroa;Apis(host) | resistance/target-site;biochemical/functional;transcriptomic |
| 10.1371/journal.pone.0031051 | 2012 | ¤a | CYP_P450;CCE_esterase;Detox_general | Varroa;Apis(host) | descriptive |
| 10.1016/j.ecoenv.2025.118723 | 2025 | Golob | AChE;GST;CCE_esterase | Varroa;Apis(host) | descriptive |
| 10.1186/s13071-016-1708-x | 2016 | Koh-Tan | GST;CYP_P450;ABC_transporter;CCE_esterase;Octopamine_R | Ixodes;other-tick | transcriptomic |
| 10.1016/j.pestbp.2025.106616 | 2025 | Wang | GST;CYP_P450;Detox_general | Varroa;Apis(host) | descriptive |
| 10.1007/s10493-025-01071-1 | 2025 | Shen | ABC_transporter | Varroa;Apis(host) | transcriptomic |
| 10.3390/insects15060409 | 2024 | Sagona | GST;Detox_general | Varroa;Apis(host) | descriptive |
| 10.1007/s10493-013-9754-y | 2014 | Dmitryjuk | AChE;CYP_P450;CCE_esterase | Varroa;other-tick;Apis(host) | descriptive |
| 10.3389/fphys.2019.01008 | 2019 | Xiong | — | other-tick | descriptive |
| 10.1016/j.vetpar.2020.109155 | 2020 | Wang | GST;CYP_P450;CCE_esterase | Dermanyssus | transcriptomic |
| 10.21203/rs.3.rs-49223/v1 | 2020 | Zhang | AChE;GST;CYP_P450;ABC_transporter;CCE_esterase | Varroa;Ixodes;other-tick | descriptive |
| 10.1007/s10519-017-9834-6 | 2017 | Hamiduzzaman | CYP_P450 | Varroa;Apis(host) | descriptive |
| 10.1016/j.jip.2017.08.005 | 2017 | O'Neal | Octopamine_R;Cl_channel | Varroa;Apis(host) | descriptive |
| 10.1007/s11356-018-3205-6 | 2018 | Hanan | — | Apis(host) | toxicology |
| 10.1016/j.pestbp.2024.105960 | 2024 | a | GST;CYP_P450;ABC_transporter;CCE_esterase | Dermanyssus | RNAi/knockdown;transcriptomic |
| 10.1016/j.vetpar.2023.109957 | 2023 | Schiavone | AChE;GST;CYP_P450;ABC_transporter;CCE_esterase | Dermanyssus;other-tick | resistance/target-site;transcriptomic |
| 10.1016/j.ibmb.2018.02.002 | 2018 | Perner | GST;ABC_transporter;Detox_general | Varroa;Dermanyssus;Ixodes;other-tick | RNAi/knockdown;transcriptomic |
| 10.1016/j.pestbp.2021.104985 | 2022 | Koç | AChE;GST;CYP_P450;CCE_esterase | Varroa;Dermanyssus | biochemical/functional |
| 10.1007/s10493-009-9249-z | 2009 | — | AChE;CCE_esterase | Dermanyssus | biochemical/functional |
| 10.1016/j.psj.2024.103612 | 2024 | Zhang | GST;CYP_P450;CCE_esterase | Dermanyssus;other-tick | RNAi/knockdown;biochemical/functional |
| 10.1093/jme/tjv252 | 2015 | Tuckow | AChE;CCE_esterase | Ixodes;other-tick | descriptive |
| 10.1016/j.sjbs.2017.01.052 | 2017 | a | GST;Detox_general | Varroa;Apis(host) | biochemical/functional |
| 10.1016/j.pestbp.2025.106364 | 2025 | Mavridis | CYP_P450;Octopamine_R | Varroa;Apis(host) | RNAi/knockdown;resistance/target-site;biochemical/functional;transcriptomic |
| 10.1016/j.pestbp.2021.104985 | 2022 | Koç | AChE;GST;CYP_P450;CCE_esterase | Varroa;Dermanyssus | descriptive |
| 10.1073/pnas.2020380118 | 2021 | Vlogiannitisa□ | AChE;GST;CYP_P450;ABC_transporter | Varroa;other-tick;Apis(host) | RNAi/knockdown;transcriptomic |
| 10.1186/1756-3305-3-73 | 2010 | (1) | — | Varroa;Apis(host) | RNAi/knockdown |
|  | 2009 | Johnson | GST;CYP_P450;CCE_esterase | Varroa;Apis(host) | toxicology |
| 10.1186/s13071-018-3020-4 | 2018 | Mangia | ABC_transporter | Ixodes;other-tick | descriptive |

| doi | year | first_ author | families | organisms | evidence |
|---|---|---|---|---|---|
| 10.1186/s13071-025-07218-8 | 2026 | Liu | CYP_P450 | Dermanyssus | biochemical/functional |
| 10.1007/s10493-023-00879-z | 2024 | Erdem | AChE;CYP_P450;CCE_esterase | Varroa;Apis(host) | resistance/target-site |
| 10.1101/2021.08.06.455389 | 2021 | Hernández-Rodríguez | AChE;CYP_P450;CCE_esterase;Octopamine_R | Varroa;Tropilaelaps;Ixodes;other-tick ;Apis(host) | resistance/target-site;transcriptomic |
| 10.3390/insects11090637 | 2020 | Siebert | CYP_P450;Detox_general | Varroa;Apis(host) | transcriptomic |
| 10.1016/j.envres.2025.123531 | 2025 | Wang | AChE;GST;CYP_P450;CCE_esterase;Detox_general | Varroa;Apis(host) | transcriptomic;toxicology |
| 10.1016/j.cbi.2012.09.010 | 2013 | Temeyer | AChE;CCE_esterase | Ixodes;other-tick | RNAi/knockdown |
| 10.1016/j.envint.2019.105256 | 2019 | Carnesecchi | AChE;GST;CYP_P450;ABC_transporter;CCE_esterase;Cl_channel;Detox_general | Varroa;Apis(host) | toxicology |
| 10.1016/j.pestbp.2020.104724 | 2020 | Wang | GST;CYP_P450 | Dermanyssus | biochemical/functional |
| 10.1073/pnas.1109535108 | 2011 | Mao | GST;CYP_P450;CCE_esterase;Detox_general | Varroa;Apis(host) | biochemical/functional |
| 10.1038/s41598-023-38187-6 | 2023 | Dawdani | AChE;CYP_P450;CCE_esterase | Varroa;Apis(host) | descriptive |
| 10.1016/j.pestbp.2025.106387 | 2025 | a | CYP_P450;Octopamine_R | Varroa;Apis(host) | resistance/target-site |
| 10.1186/s12870-025-06108-6 | 2025 | Sakla | GST | Varroa;Apis(host) | descriptive |
| 10.1016/j.jinsphys.2016.03.004 | 2016 | Chaimanee | CYP_P450;Detox_general | Varroa;Apis(host) | transcriptomic |
| 10.1186/s13071-016-1497-2 | 2016 | Mangia | ABC_transporter | Ixodes;other-tick | transcriptomic |
| 10.1371/journal.pone.0054092 | 2013 | Johnson | CYP_P450;CCE_esterase;Octopamine_R | Varroa;Apis(host) | biochemical/functional |

### Table S16. Phase-2 curated literature evidence

Curated evidence supporting the receptor, channel and signalling target families. *obtained* records whether the full text was consulted.

| doi | family_axis | organism | evidence_type | key_finding_paraphrase | year | obtained |
|---|---|---|---|---|---|---|
| 10.7554/elife.68268 | GPCR_aminergic / Octβ2R | Varroa+Apis | functional·selectivity | Amitraz & metabolite DPMF activate all 4 Varroa octopamine receptors (nM EC50); Octβ2R is the sole in vivo target; honeybee Octβ2R 16x less amitraz-sensitive via 3 bee-specific residues (E208V/I335T/I350V) — molecular basis of the bee-vs-mite selectivity window. | 2021 | yes |
| 10.1016/j.pestbp.2024.106080 | GPCR_aminergic / Octβ2R | Varroa | functional·CRISPR | CRISPR-Cas9 validation of a novel β-adrenergic-like octopamine-receptor mutation associated with amitraz resistance. | 2024 | yes |
| 10.1038/s41598-025-85279-6 | GPCR_aminergic / Octβ2R + VGSC | Varroa | field·genotype-phenotype | Alberta: Octβ2R Y215H in 90% of apiaries correlates with phenotypic amitraz resistance; VGSC L925I/M in 100% of apiaries (kdr). | 2025 | yes |
| 10.21203/rs.3.rs-9276822/v1 | GPCR_aminergic / Octβ2R | Varroa | field·genotype | France 2018-25: Octβ2R N87S the only amitraz mutation detected; RR frequency rises with resistance class (0.03→0.39); resistance persists after amitraz withdrawal. | 2026 | yes |
| 10.1016/j.rvsc.2026.106101 | GPCR_aminergic / Octβ2R | Varroa | field·caution | Central Spain: Octβ2R F290L at 89% (all homozygous) independent of amitraz exposure — historical standing variation, NOT a selection signal; cautions against treating every Octβ2R SNP as a resistance marker. | 2026 | yes |
| 10.1371/journal.pone.0082941 | VGSC_Na (para) | Varroa | molecular·diagnostic | Landmark L925V in VGSC IIS5: 100% of tau-fluvalinate survivors carry it; <10% in untreated; TaqMan assay developed. | 2013 | yes |
| 10.1016/j.ibmb.2006.08.006 | VGSC_Na (para) | Varroa(heterologous) | functional | Varroa L1770P (cockroach P1577L equivalent) expressed in BgNav increases fluvalinate sensitivity 5-fold without altering gating; native proline in insect channels underlies their lower pyrethroid sensitivity (arachnid-vs-insect selectivity). | 2006 | yes |
| 10.1371/journal.pone.0155332 | VGSC_Na (para) | Varroa | molecular·genotype-phenotype | US: novel L925M & L925I; 98% of tau-fluvalinate survivors carry a mutant allele; survival vs M925/I925 correlation ρ=0.997. | 2016 | yes |
| 10.1007/s10493-021-00665-9 | VGSC_Na (para) | Varroa | molecular·resistance-ratio | Belgium: L925V dominant (38-44%); resistant population shows 12.6-fold flumethrin resistance ratio (LC50). | 2021 | yes |
| 10.3390/insects16060548 | VGSC_Na (para) | Varroa | field·genotype-phenotype | Muğla, Türkiye: phenotypic + genotypic flumethrin resistance assessment in Varroa. | 2025 | yes |

| doi | family_axis | organism | evidence_type | key_finding_paraphrase | year | obtained |
|---|---|---|---|---|---|---|
| 10.1007/s10493-026-01132-z | VGSC_Na (para) | Varroa | field·genotype | Türkiye: widespread VGSC codon-925 kdr resistance across 10 provinces (PCR-RFLP). | 2026 | yes |
| 10.1007/s10493-025-01002-0 | VGSC_Na (para) | Varroa | field·genotype | Türkiye beekeeping areas: status/frequencies of VGSC kdr mutations. | 2025 | yes |
| 10.1007/s10493-026-01121-2 | VGSC_Na (para) | Varroa | method·diagnostic | Snapback HRM assay to detect/differentiate L925 kdr mutations rapidly. | 2026 | yes |
| 10.1016/j.pestbp.2023.105655 | VGSC_Na (para) | Varroa | method·monitoring | Korea: combined bioassay + molecular-marker monitoring of fluvalinate resistance. | 2023 | yes |
| 10.1002/ps.7126 | VGSC_Na (para) | Varroa | molecular·phenotype-genotype | Relationship between in-vitro tau-fluvalinate phenotype and VGSC L925V genotype. | 2022 | yes |
| 10.1016/j.ibmb.2014.10.002 | LGIC_GABA_RDL | Varroa | functional·pharmacology | Varroa RDL M2 carries 4 atypical residues; T6'M ablates picrotoxin block and converts thymol from potentiator to inhibitor; lowers GABA EC50 — channel-level basis for differential pharmacology. | 2014 | yes |
| 10.1074/jbc.ra118.005365 | LGIC_GABA_RDL | Varroa | functional·repertoire | Six GABA-channel subunits cloned (VdesRDL1-4, LCCH3, GRD); RDL1-3 form functional homomers, GRD/LCCH3 a cationic heteromer; fipronil IC50 spans 0.9-55 μM; VdesRDL1 fipronil-resistance via S2'/M6'. | 2018 | yes |
| 10.1016/j.pestbp.2022.105064 | LGIC_GABA_RDL | Varroa+Apis | functional·selectivity | Varroa 'B' RDL (T6'M) is inhibited by thymol whereas Apis RDL is potentiated — T6'M drives the Varroa-vs-Apis difference in thymol action. | 2022 | yes |
| 10.1186/s12864-022-08669-4 | cys-loop LGIC (comparative) | Ixodes ricinus | comparative·transcriptome | Synganglion meta-transcriptome (96.6% BUSCO); 46 non-AChR cys-loop genes; 8 nAChR clades (β2/α5 lack insect homologs); GABA-1/Rdl with exon-triplication isoforms; GluCls resolved. | 2022 | yes |
| 10.1101/2021.12.20.473502 | cys-loop LGIC (comparative) | Ixodes ricinus | comparative·functional | Companion synganglion study; I. ricinus RDL functional in oocytes (GABA EC50 ≈20 μM); full cys-loop LGIC phylogeny (GABA, pHCl, GluCl, nAChR clades). | 2021 | yes |
| 10.1371/journal.pone.0102667 | Neuropeptide/GPCR + NR (comparative) | Ixodes scapularis | comparative·transcriptome | Female synganglion: 15 neuropeptides, 14 neuropeptide receptors, 6 aminergic/transmitter receptors (ACh/GABA/dopamine/glutamate/octopamine/serotonin) and an ecdysone nuclear receptor expressed — confirms the Phase-2 GPCR/NR/channel repertoire in the Acari CNS. | 2014 | yes |
| 10.1038/s44386-026-00050-9 | VGSC / channel-acting toxin | Varroa+Apis | functional·MoA·safety | Spider-venom peptides Ht1a/Gg1a are varroacidal; sHt1a inhibits both Varroa and Apis NaV ~25% (no channel-level selectivity) yet is mite-selective in vivo (safe to bees at 52x dose); inactive on human NaV/CaV/nAChR. | 2026 | yes |
| 10.17420/ap71.541 | Comparative target / reproduction | Dermanyssus gallinae | review·RNAi | D. gallinae biology/genetics review; RNAi of vitellogenin and TOR impairs reproduction — cross-links to the Phase-3 TOR/reproduction axis. | 2025 | yes |

### Table S17. Bidirectional over-call and under-call across 31 homology criteria

Both error directions recomputed from one search output over a grid of identity and coverage criteria, on 2,210 group-absent and 5,336 group-present Acari-conserved genes. *E1e-3* is the E-value criterion reported in the main text, which imposes no identity or coverage floor. *ratio* is over-call divided by under-call.

| criterion | min_identity | min_coverage | n_og_absent | n_overcalled | overcall_pct | n_og_present | n_undercalled | undercall_pct | ratio |
|---|---|---|---|---|---|---|---|---|---|
| E1e-3 | | | 2210 | 1489 | 67.38 | 5336 | 67 | 1.26 | 53.66 |
| id20_scov20 | 20 | 20 | 2210 | 1599 | 72.35 | 5336 | 122 | 2.29 | 31.65 |
| id20_scov30 | 20 | 30 | 2210 | 1466 | 66.33 | 5336 | 203 | 3.80 | 17.44 |
| id20_scov40 | 20 | 40 | 2210 | 1329 | 60.14 | 5336 | 349 | 6.54 | 9.19 |
| id20_scov50 | 20 | 50 | 2210 | 1208 | 54.66 | 5336 | 523 | 9.80 | 5.58 |
| id20_scov70 | 20 | 70 | 2210 | 947 | 42.85 | 5336 | 975 | 18.27 | 2.35 |
| id25_scov20 | 25 | 20 | 2210 | 1470 | 66.52 | 5336 | 207 | 3.88 | 17.15 |
| id25_scov30 | 25 | 30 | 2210 | 1333 | 60.32 | 5336 | 313 | 5.87 | 10.28 |
| id25_scov40 | 25 | 40 | 2210 | 1198 | 54.21 | 5336 | 474 | 8.88 | 6.10 |
| id25_scov50 | 25 | 50 | 2210 | 1081 | 48.91 | 5336 | 657 | 12.31 | 3.97 |
| id25_scov70 | 25 | 70 | 2210 | 842 | 38.10 | 5336 | 1106 | 20.73 | 1.84 |
| id30_scov20 | 30 | 20 | 2210 | 1143 | 51.72 | 5336 | 615 | 11.53 | 4.49 |
| id30_scov30 | 30 | 30 | 2210 | 1039 | 47.01 | 5336 | 747 | 14.00 | 3.36 |

| criterion | min_identity | min_coverage | n_og_absent | n_overcalled | overcall_pct | n_og_present | n_undercalled | undercall_pct | ratio |
|---|---|---|---|---|---|---|---|---|---|
| id30_scov40 | 30 | 40 | 2210 | 929 | 42.04 | 5336 | 898 | 16.83 | 2.50 |
| id30_scov50 | 30 | 50 | 2210 | 833 | 37.69 | 5336 | 1066 | 19.98 | 1.89 |
| id30_scov70 | 30 | 70 | 2210 | 642 | 29.05 | 5336 | 1498 | 28.07 | 1.03 |
| id35_scov20 | 35 | 20 | 2210 | 794 | 35.93 | 5336 | 1263 | 23.67 | 1.52 |
| id35_scov30 | 35 | 30 | 2210 | 722 | 32.67 | 5336 | 1376 | 25.79 | 1.27 |
| id35_scov40 | 35 | 40 | 2210 | 626 | 28.33 | 5336 | 1511 | 28.32 | 1.00 |
| id35_scov50 | 35 | 50 | 2210 | 549 | 24.84 | 5336 | 1656 | 31.03 | 0.80 |
| id35_scov70 | 35 | 70 | 2210 | 434 | 19.64 | 5336 | 2039 | 38.21 | 0.51 |
| id40_scov20 | 40 | 20 | 2210 | 569 | 25.75 | 5336 | 1906 | 35.72 | 0.72 |
| id40_scov30 | 40 | 30 | 2210 | 504 | 22.81 | 5336 | 2006 | 37.59 | 0.61 |
| id40_scov40 | 40 | 40 | 2210 | 425 | 19.23 | 5336 | 2124 | 39.81 | 0.48 |
| id40_scov50 | 40 | 50 | 2210 | 381 | 17.24 | 5336 | 2257 | 42.30 | 0.41 |
| id40_scov70 | 40 | 70 | 2210 | 302 | 13.67 | 5336 | 2576 | 48.28 | 0.28 |
| id50_scov20 | 50 | 20 | 2210 | 246 | 11.13 | 5336 | 3272 | 61.32 | 0.18 |
| id50_scov30 | 50 | 30 | 2210 | 213 | 9.64 | 5336 | 3347 | 62.72 | 0.15 |
| id50_scov40 | 50 | 40 | 2210 | 189 | 8.55 | 5336 | 3428 | 64.24 | 0.13 |
| id50_scov50 | 50 | 50 | 2210 | 166 | 7.51 | 5336 | 3516 | 65.89 | 0.11 |
| id50_scov70 | 50 | 70 | 2210 | 144 | 6.52 | 5336 | 3684 | 69.04 | 0.09 |
